# Lipid-dependent conformational dynamics of bacterial ATP-binding cassette transporter Sav1866

**DOI:** 10.1101/2024.04.18.590185

**Authors:** Shadi A Badiee, Jeevapani Hettige, Mahmoud Moradi

## Abstract

Sav1866, a bacterial ATP-binding cassette (ABC) exporter, plays a crucial role in cellular processes by facilitating the efflux of a diverse range of substrates, including drugs, chemotherapeutic agents, peptides, and lipids. This efflux activity significantly impacts the effectiveness of various therapies against bacterial infections. In our recent investigation, we focused on understanding the conformational dynamics of Sav1866 within different lipid environments. Specifically, we explored its behavior in environments composed of DMPC and POPE lipids, which exhibit crucial distinctions not only in their headgroup polarity but also in the length and saturation of their hydrophobic tails. Our extensive set of equilibrium microsecond-level all-atom molecular dynamics (MD) simulations revealed significant distinctions in transporter behavior influenced by these lipid compositions. We observed a rapid transition to an occluded-inward-facing (IF-occ) conformation in POPE environments, contrasting with a channel-like behavior in DMPC environments, deviating from the expected alternating access mechanism (AAM). These findings underscore the significant impact of lipid compositions on ABC transporter function, offering new perspectives on membrane transport mechanisms.

## INTRODUCTION

The ATP-binding cassette (ABC) transporter superfamily stands as one of the most extensive membrane protein families, present across a wide range of organisms, ranging from prokaryotes to humans [2]. ABC transporters harness the energy derived from the binding and subsequent break-down of ATP molecules to transport a variety of substrates across the membrane. The nucleotide-binding domain (NBD) and the transmembrane domain (TMD) are the two primary components of ABC transporters. The transmembrane domain (TMD) creates a pathway for transporting substances by comprising twelve transmembrane (TM) helices and substrate-binding site. However, NBD is the site where ATP binds and gets hydrolyzed [3, 4, 45]. Membrane transporters efficiently transport their substrates through significant structural changes, alternating between inward-facing (IF) and outward-facing (OF) states in a process known as the alternating access mechanism [5]. In this mechanism, the protein’s substrate binding site is exposed alternately inside (IF) or outside (OF) the membrane (IF *↔* OF), potentially going through a number of intermediate phases so that ATP binding triggers NBD dimerization and the OF state’s development. ADP and Pi are released due to ATP hydrolysis, which is how the NBD dissociates, and the IF state is restored [30, 31, 47].

Membrane transporters, particularly those in the ABC superfamily, are known to be greatly influenced by the lipid content of their membranes [1, 35–37, 41, 42, 46]. Our study focuses on a bacterial multidrug ABC exporter named Sav1866, which exists as a homodimer. This homodimeric structure comprises two Transmembrane Domains (TMDs) and two Nucleotide-Binding Domains (NBDs) [48]. Sav1866 has been evidenced to facilitate the transportation of amphiphilic compounds across cellular membranes [49, 50]. In our earlier investigation [1], our research team investigated the behavior of the Sav1866 when exposed to various lipids, with a particular emphasis on the roles of PC and PE lipids. This study involved simulations lasting 2.4 *µ*s each, which provided valuable insights into how the lipid environment influences the protein’s behavior.

In our current study, we are advancing our research by significantly extending the simulation duration to 30 *µ*s for POPE and performing 10 *µ*s simulations in DMPC, which was not part of our previous work. This extension enables us to investigate the distinct effects of DMPC and compare them with POPE, two lipid types that exhibit differences not only in their headgroups but also in their tail structures. This extensive study attempts to provide a more comprehensive understanding of how these proteins function in various lipid environments.

Molecular dynamics simulations are known to be a valuable tool for detailed atomic-level studies of proteins and membrane transporters [32–34]. Numerous MD simulations have been conducted to investigate the dynamics of various ABC transporters [5–7, 9, 10, 32, 34, 39, 47]. To our knowledge, this study represents the first comprehensive computational exploration into how the presence of DMPC lipids influences Sav1866’s alternating access mechanism. What sets our research apart is the remarkable length of our all-atom MD simulations in both lipid compositions (POPE and DMPC), which provides an extensive and detailed perspective on the subject.

## RESULTS

The nucleotide-free apo model of Sav1866 was created by extracting the nucleotides from the crystal structure of the nucleotide-bound outward-facing (OF) state of Sav1866 (PDB: 2HYD) [11, 48]. This apo model was then incorporated into two distinct lipid environments, such as POPE and DMPC lipid bilayers. We conducted significantly extended simulations, lasting 30 *µ*s and 10 *µ*s for POPE and DMPC, respectively. These durations qualify as substantially long-lasting MD simulations, enabling a comprehensive system exploration. We noted a remarkable contrast in the behavior of the ATP-removed outward-facing (OF) state when placed in the POPE and DMPC lipid environments. In particular, the protein in the POPE environment exhibited behavior consistent with the alternating access mechanism, aligning with expectations. In contrast, the protein within the DMPC environment demonstrated an unexpected channel-like behavior, which is inconsistent with the traditional alternating access mechanism. To streamline our discussion, we initiate with an examination of the initial 10 *µ*s of the POPE and DMPC simulations, revealing significant differences in their behavior. Subsequently, we conduct an in-depth analysis of the 30 *µ*s POPE simulation. Additionally, our study uncovers interesting side stories, such as evidence of Sav1866’s flippase activity within DMPC simulation.

### Sav1866 Elicits a Channel-like Conformation in DMPC, While POPE Induces an IFocc State

We initiate a comprehensive root mean square deviation (RMSD) analysis to elucidate the structural dynamics of Sav1866 in two distinct lipid environments, namely POPE and DMPC. The RMSD plot (Figure. 1A) illustrates the deviation of C*α* atoms of the protein with respect to the initial model. This analysis reveals an increase in the RMSD value of the protein in both lipid environments, ranging from 2 to 6 Å throughout the initial 5 *µ*s simulation. After the initial 5 *µ*s, the RMSD in the POPE environment stabilizes below 6 Å, maintaining a relatively constant behavior. However, the protein in the DMPC environment exhibits a different pattern, with its RMSD peaking at 8 Å before converging back to around 6 Å. We further measured the internal RMSD for both the Transmembrane Domain (TMD) and the Nucleotide-Binding Domain (NBD) in both lipid compositions (Figure. S1).The NBD domain consistently exhibits higher RMSD values than the TMD. This difference may be attributed to ATP segregation, rendering the TMD more stable compared to the NBD. To summarise our findings from the RMSD analyses, during the initial 5 *µ*s, the protein behaves similarly in both lipid compositions. However, as the simulation progresses, they start to behave differently, which indicates how much the lipid content affects the structural dynamics of Sav1866.

**FIG. 1.**
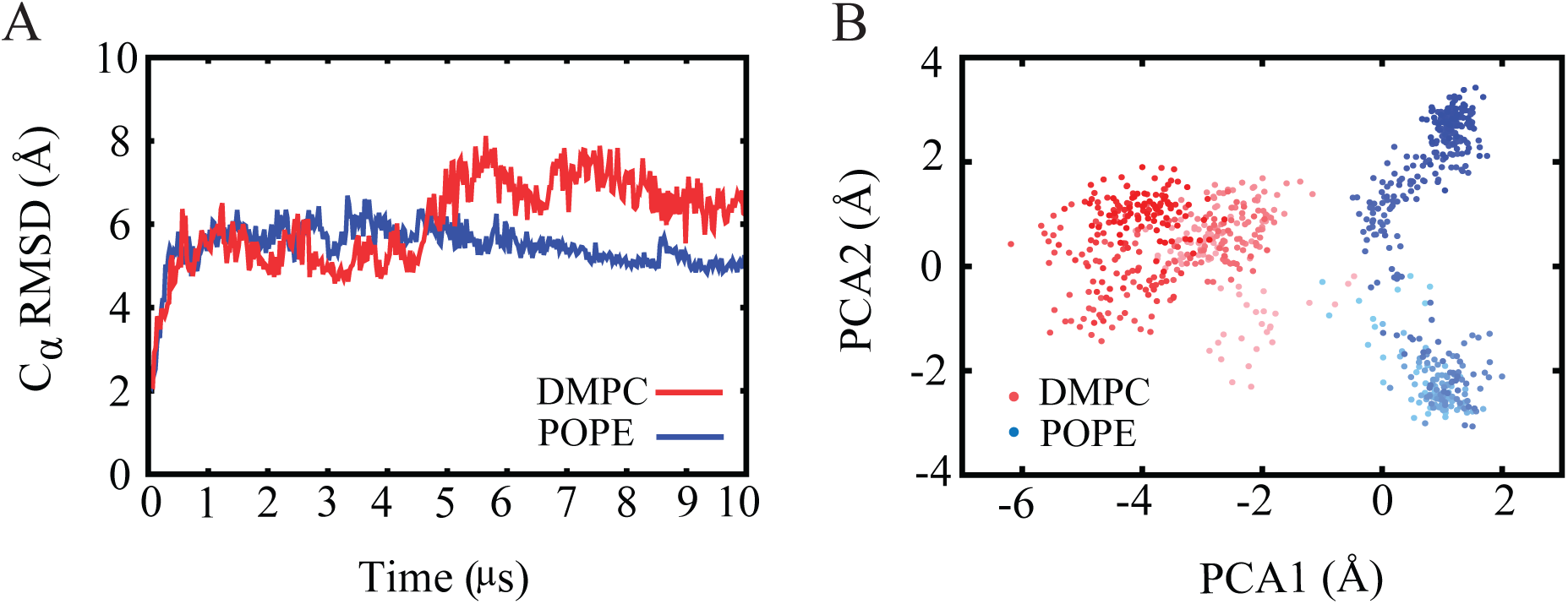
(A) Time series of root mean square deviation (RMSD) of C*α* atoms of the protein associated with apo protein in POPE, DMPC with respect to the initial model. (B) Projection of POPE and DMPC trajectories onto the (PC1, PC2) space, which represents the first two principal components of protein C*α* atoms.

We also conducted three independent repeats of molecular dynamics simulations for Sav1866 embedded in lipid bilayers composed of DMPC, POPC, DOPC, and DPPC, each lasting 2.4 *µ*s. The calculated root mean square deviation (RMSD) of C*α* atoms for the protein was analyzed, and the data is provided in the supporting information (Figure. S2). Even within the relatively short duration of these simulations, a distinct trend in the stability of Sav1866 has been observed. For instance, the RMSD values indicate that Sav1866 experiences less stability in DMPC (with 14 carbons in each chain and saturated) lipid composition, with the largest RMSD value observed. Additionally, DPPC with 16 carbons in each saturated chain exhibits lower RMSD, followed by DOPC with two unsaturated chains of 18 carbons. Interestingly, the RMSD values for the last DOPC and POPC (chains including 16 and 18 carbons and one unsaturated bond) simulations are relatively close despite variations in chain length and unsaturation. The observed RMSD trends suggest a correlation between lipid composition, chain size, and saturation levels with the stability of Sav1866. DMPC, with shorter and saturated chains, results in less stability of the protein, whereas DOPC and POPC with longer and unsaturated chains exhibit increased stability. By increasing the chain size, the stability of the protein is also increased, but at the same time, saturation or unsaturation has effects on it. The longer chain size and the presence of unsaturated bonds in DOPC and POPC appear to contribute to a stabilization effect. Furthermore, internal RMSD calculations for the Nucleotide Binding Domain (NBD) and Transmembrane Domain (TMD) also revealed interesting insights. The NBD displays a lower stability compared to TMD, as evidenced by higher RMSD values, which are detailed in the supporting information (Figure. S3).

In order to compare the global protein conformational changes in the two environments, we have performed principal component analysis (PCA) using a combination of both POPE and DMPC trajectories using 10 *µ*s of each. When projected onto the first two principal components (PC1 and PC2), it is evident that the transporter occupies different regions of the conformational landscape in the POPE and DMPC environments (Figure. 1B). The PCA plot, which features a color gradient from light to dark, illustrates the transitional dynamics within both systems. In both the POPE and DMPC environments, we observe the transitions occurring within a similar time-frame, somewhere in the middle of the simulation. Additionally, the plot for POPE reveals two distinct populations that appear reasonably well-separated. Initially, it drops to the lower minimum and subsequently shifts to the upper minimum, indicating significant conformational changes. In contrast, the DMPC plot suggests an initial occupancy on the right side, with a migration towards the left side at a certain point in the simulation. This analysis provides a comprehensive understanding of the distinct behavior exhibited by Sav1866 in response to these lipid environments.

To gain deeper insights into the global protein conformational changes linked to the opening and closing of the cytoplasmic and periplasmic sides of the protein, we measured the angles between the two bundles of transmembrane (TM) helices in the inward-facing (IF) state and the outward-facing (OF) state, known as the *α* and *β* angle, respectively (Figure. 2). If we consider the protein in its inward-facing (IF) state, it consists of two bundles (Figure. 2A). The first bundle comprises TM helices 1, 2, 3, and 6 from monomer A, along with TM helices 4 and 5 from monomer B. The second bundle is composed of TM helices 1, 2, 3, and 6 from monomer B and TM helices 4 and 5 from monomer A. In the data, an *α* angle below about 35*^◦^* (Figure. 3A) suggests the cytoplasmic gate closes within the initial 1 *µ*s in the POPE environment and remains closed throughout the simulation. Alternatively, in the DMPC environment, the angle shows fluctuations and an overall increase from 30*^◦^* to 50*^◦^*.

**FIG. 2.**
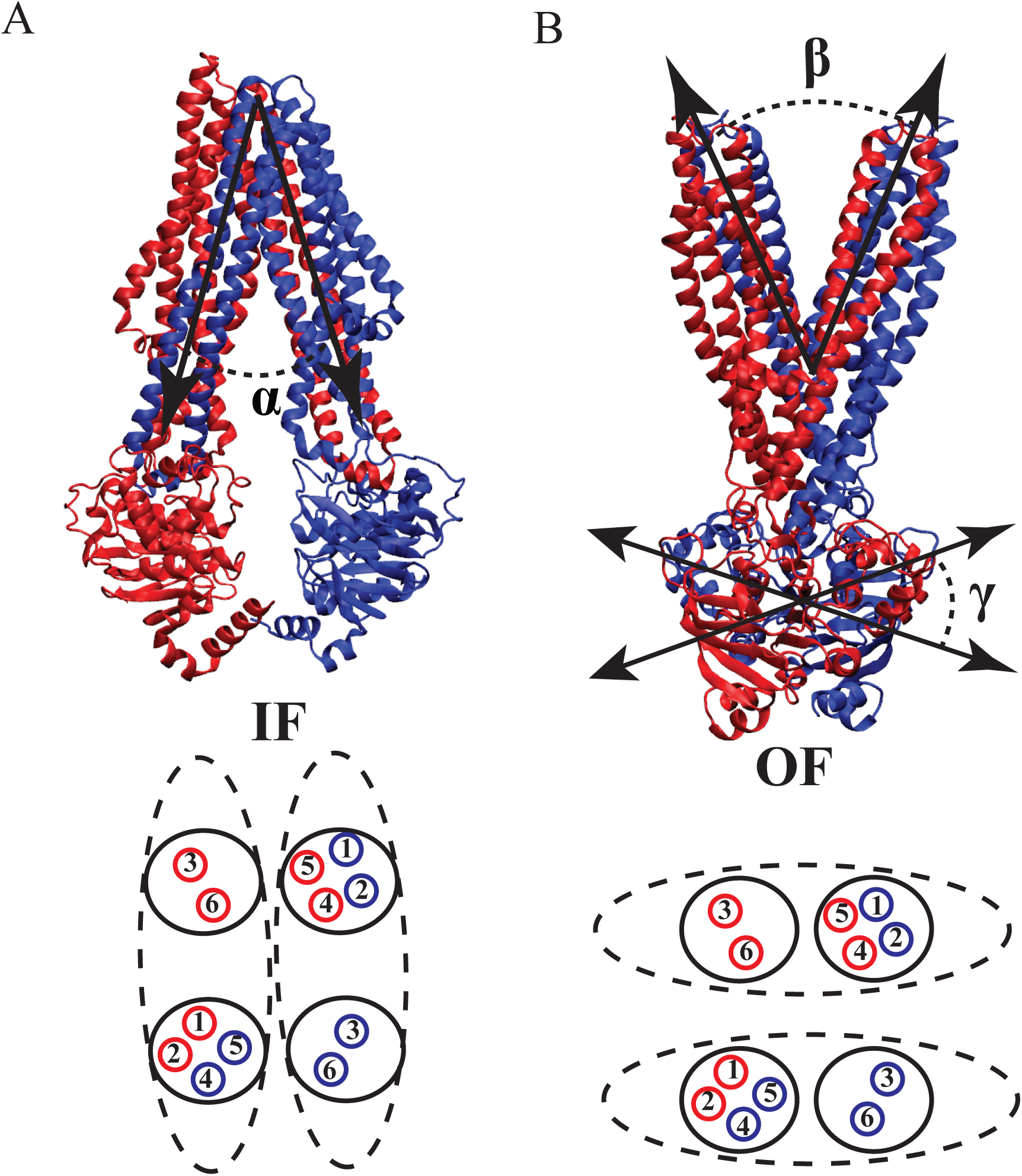
The reaction coordinates *α*, *β*, and *γ*. (A)In the IF conformation, the *α* angle is the angle between two bundles, with each bundle comprising TM helix numbers 1, 2, 3, and 6 from one monomer and helix 4 and 5 from another monomer. The *β* angle represents the angle between two bundles in the OF conformation. (B) In the OF conformation, each bundle includes TM helix numbers 1 and 2 from one monomer and 3, 4, 5, and 6 from another monomer. The *γ* angle represents the angle between two NBDs.

**FIG. 3.**
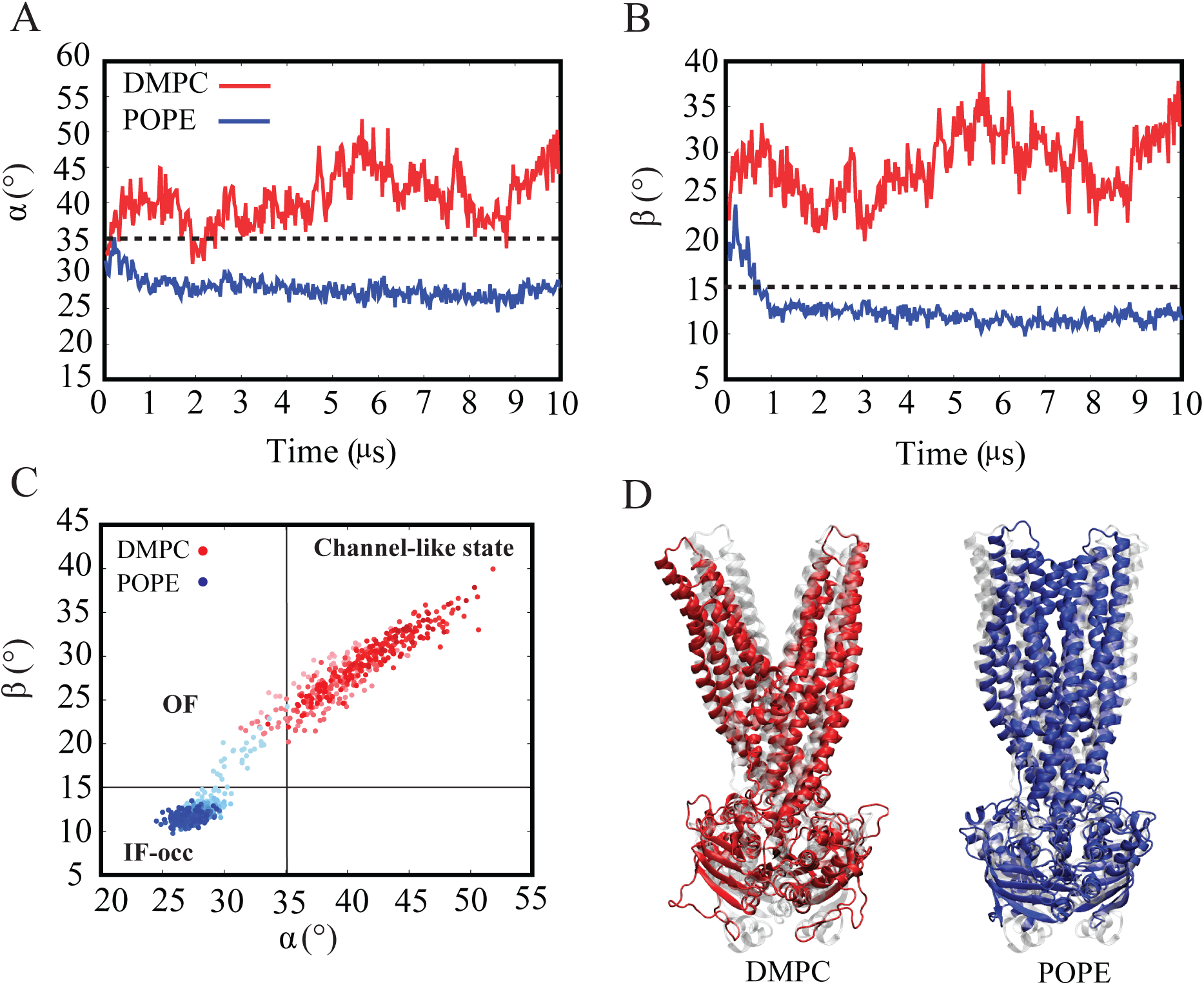
(A) Time series of the *α* angle (cytoplasmic gate) and (B) *β* angle (periplasmic gate) in the presence of both POPE and DMPC. The POPE data is represented in blue, while the DMPC data is shown in red. (C) display the outcome of projecting the alpha and beta angles in relation to each other. The MD simulations conducted within a DMPC lipid environment result in conformations that are accessible from both the cytoplasmic and periplasmic sides (red color), in contrast to the expected alternating access mechanism. The data clearly illustrates the transition from the OF to the IF-occ state when Sav1866 is embedded in a POPE lipid composition (blue color). (D) Molecular images of the protein in the last frame of the simulation. The protein is shown embedded in DMPC (red) and POPE (blue), with its initial frame depicted in transparent gray for comparison.

On the other hand, the *β* angle (Figure. 3B), characterizing the periplasmic side of the protein in the outward-facing (OF) state, provides insight into the global protein conformational changes. We employ a 15*^◦^* threshold to assess the closure of the Transmembrane Domains (TMDs) on the periplasmic side. In the POPE lipid environment, the plot indicates the closure occurring within the initial 1 *µ*s, and this closed state persists for the entire 10 *µ*s simulation. In contrast, protein in the DMPC lipid environment does not result in the closed state, with the angle consistently exceeding 15*^◦^* throughout the entire 10 *µ*s simulation. It starts at 22*^◦^* in the first frame and reaches approximately 38*^◦^* in the final frame. The projection of the *β* angle and the *α* angle (Figure. 3C), illustrating the protein’s conformational changes in different lipid environments, indicates that in the POPE lipid environment, the protein undergoes a transition from the outward-facing (OF) state to the occluded inward-facing (IF-occ) state. In contrast, in the DMPC environment, the protein demonstrates a distinct behavior within the context of the alternating access mechanism (AAM). It deviates from the standard AAM by moving towards a state we refer to as the “channel-like state”, characterized by both gates being open, representing a significant difference from the expected behavior.

In our previous PC simulations with 2.4 *µ*s, we consistently observed the periplasmic gate opening in all cases (DMPC, DPPC, DOPC, and POPC) (Figure. S4) However, a unique behavior, resembling a channel-like structure with simultaneous opening of both cytoplasmic and periplasmic gates, was exclusively seen in the DMPC simulation. This specific feature sets DMPC apart from other PC simulations even in a short time of the simulation. In one repeat of the DPPC simulation, we noticed changes in the cytoplasmic angle, reaching values higher than 35 degrees. It’s important to note that these changes were not as frequent as observed in the DMPC simulation.

### Water Density Maps Provide Evidence of an Expanded Pore in the DMPC Lipids, Suggesting an Open Periplasmic Gate and Partially Open Cytoplasmic Gate in the Protein

Water density maps were employed to visualize the differential behavior of cytoplasmic and periplasmic gates in DMPC and its contrast with POPE. These maps effectively estimate water molecule distribution around and within the protein. We focused on the final 5 *µ*s of the 10 *µ*s simulations in both POPE and DMPC environments (Figure. 4A, B). In the case of DMPC, the water density maps reveal an interesting pattern. Although the widened pore on the cytoplasmic side does not extend fully into the cytoplasm, it indicates a partial opening toward the cytoplasm. This behavior aligns more closely with what we term a “channel-like state”. The water density map presented (Figure. 4) is based on standard deviation, and the maximum density map, showing the maximum water occupancy, is provided in the Supporting Information (Figure. S5).

**FIG. 4.**
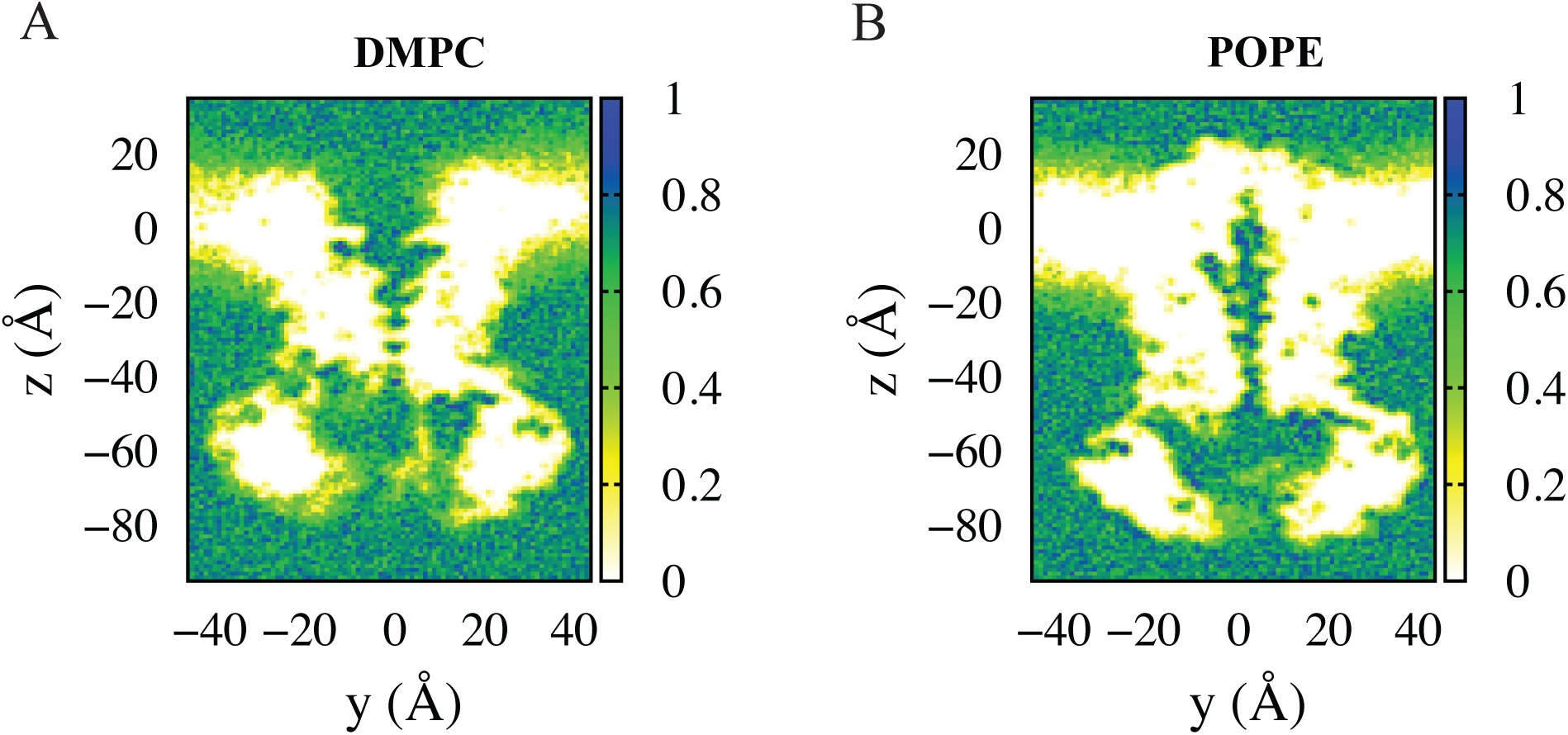
Water density, represented based on standard deviation, is depicted in and around the transporter, as observed within a 1 Å thick cross-section of the simulation box during the last 5 *µ*s of both DMPC (A) and POPE (B) simulations.

To evaluate water distribution within the pore and validate the water density map, we calculated water accessibility along the pore’s Z-axis during the final 5 *µ*s of the simulations (Figure. 5), given the distinct behavior of the protein in various lipid compositions during this period. The resulting plot clearly demonstrates that the average water count in the DMPC environment exceeds that in the POPE environment along the Z-axis. This analysis provides further evidence of Sav1866’s “channel-like state” behavior in DMPC lipid composition and the protein’s variable behavior in response to various lipid environments.

**FIG. 5.**
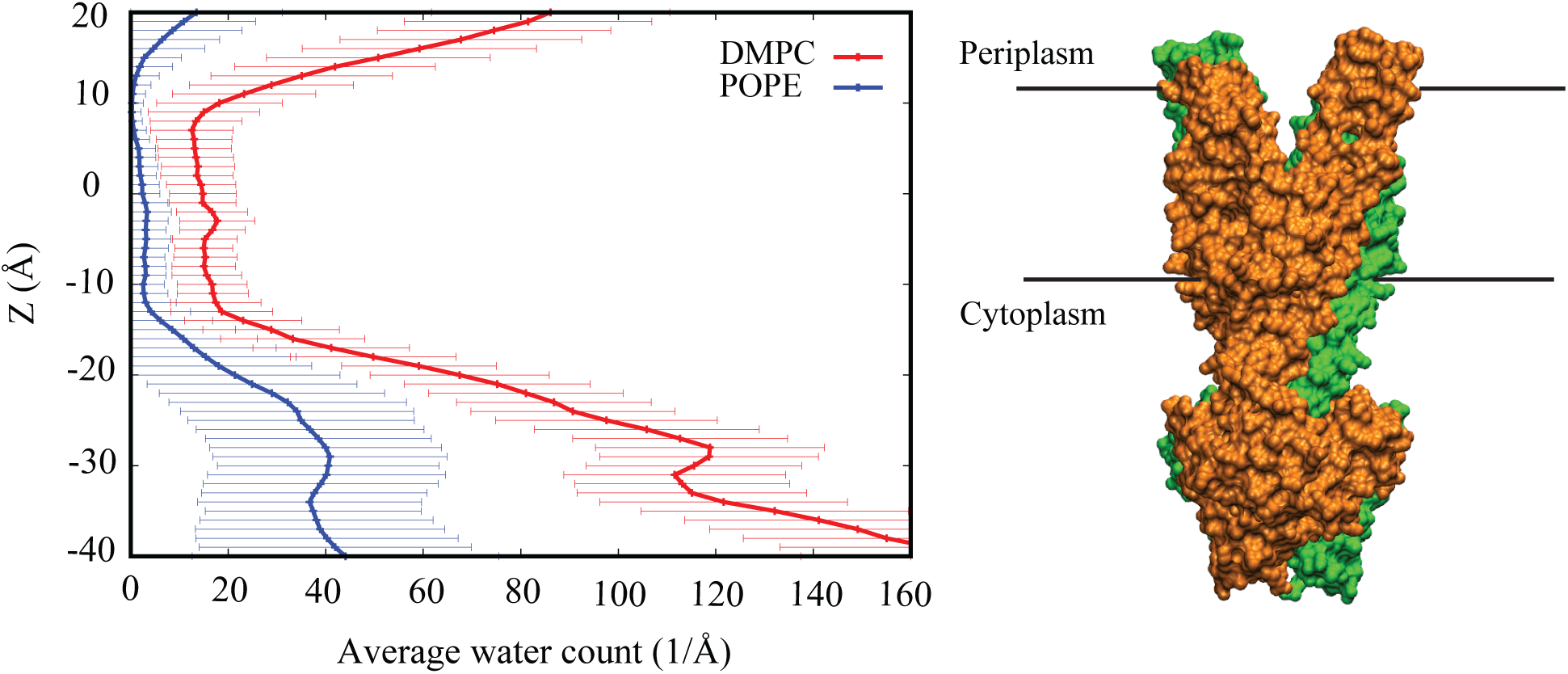
Water accessibility along the Z-axis during the final 5 *µ*s of the simulations is presented. DMPC and POPE are shown in red and blue colors, respectively. The resulting plot illustrates that the average water count in the DMPC environment is significantly higher than in the POPE environment along the Z-axis, particularly in the periplasmic and cytoplasmic gates, which are located at approximately 10 Å and -10 Å, respectively.

### Stability of TMD Hinge and the Role of Salt Bridges in Opening and Closing the Gates

In the previous study conducted by our research group [1], over a 2.4 *µ*s duration, significant secondary structural changes within one monomer (monomer A) of Sav1866 in H3 (residues 138-141) have been observed in the presence of POPE lipids. This transition led to the formation of a TMD hinge region. This structural change was accompanied by the forming of a salt bridge between R81 and D145, which was located slightly above the hinge region and close to the membrane center. They hypothesized that the newly formed salt bridge was critical in triggering the periplasmic gate closure in the POPE lipid environment.

In the current comprehensive simulations, we observed that this secondary structural transition within H3 remains stable for the whole simulation time in the POPE environment (Figure. 6A). However, during our 10 *µ*s simulation in DMPC, this transition and formation of the essential R81-D145 salt bridge were not observed. Therefore, the closure of the periplasmic gate does not happen in DMPC. Furthermore, in the POPE simulations, the development of the R81-D145 salt bridge (Figure. 6E) was associated with the weakening of the intrabundle salt bridge between R296 and D145 (Figure. 6G), which linked H6 and H3 inside the same monomer. The development of the hinge area in H3 also indicated that the formation of the R81-D145 salt bridge did not entirely remove the R296-D145 salt bridge. This feature continued in our current long-term POPE simulations, demonstrating consistent behavior (Figure. 6E, G). In contrast, our 10 *µ*s simulations in the DMPC environment revealed a completely different picture for Sav1866. Throughout the simulation, the R296-D145 salt bridge (Figure. 6F) stayed steady and stable, and the crucial R81-D145 salt bridge (Figure. 6D) never formed. These findings emphasize how differently Sav1866 behaves when exposed to various lipid compositions and how these salt bridges affect the conformational dynamics of the protein. Additionally, we also identified the presence of an intrabundle salt bridge between D127 (H3) and R313 (H6) in monomer A, located below the hinge region of H3 and adjacent to the cytoplasmic gate (Figure. 7C and D). This salt bridge, originally observed in the initial crystal structure, initially broke but later reformed and remained stable from 2.1 *µ*s until the end of the simulation in the POPE environment (Figure.7D). However, in the DMPC environment, it was disrupted and could not reform by the end of the 10 *µ*s simulation (Figure.7C). Simultaneously, the formation of the intrabundle salt bridge between D127 (H3) and R313 (H6) in monomer A of POPE (Figure.7D) led to the disruption of the interbundle salt bridge between E129 (H3) and R97 (H2) (Figure. 7B). On the opposing side, the behavior in DMPC was significantly different, as the E129 (H3)-R97 (H2) interbundle salt bridge (Figure. 7A) formed as a result of the breakdown of the D127 (H3)-R313 (H6) intrabundle salt bridge (Figure. 7C). The difference in the stability of the intrabundle salt bridge and the breakdown of the interbundle salt bridge in the POPE environment suggests a potential role in maintaining the closure of the cytoplasmic gate. However, in the DMPC environment, these observations suggest that the disruption of the intra-bundle salt bridge (Figure. 7C) and the increased stability of the interbundle salt bridge (Figure. 7A) may contribute to the opening of the cytoplasmic gate.

**FIG. 6.**
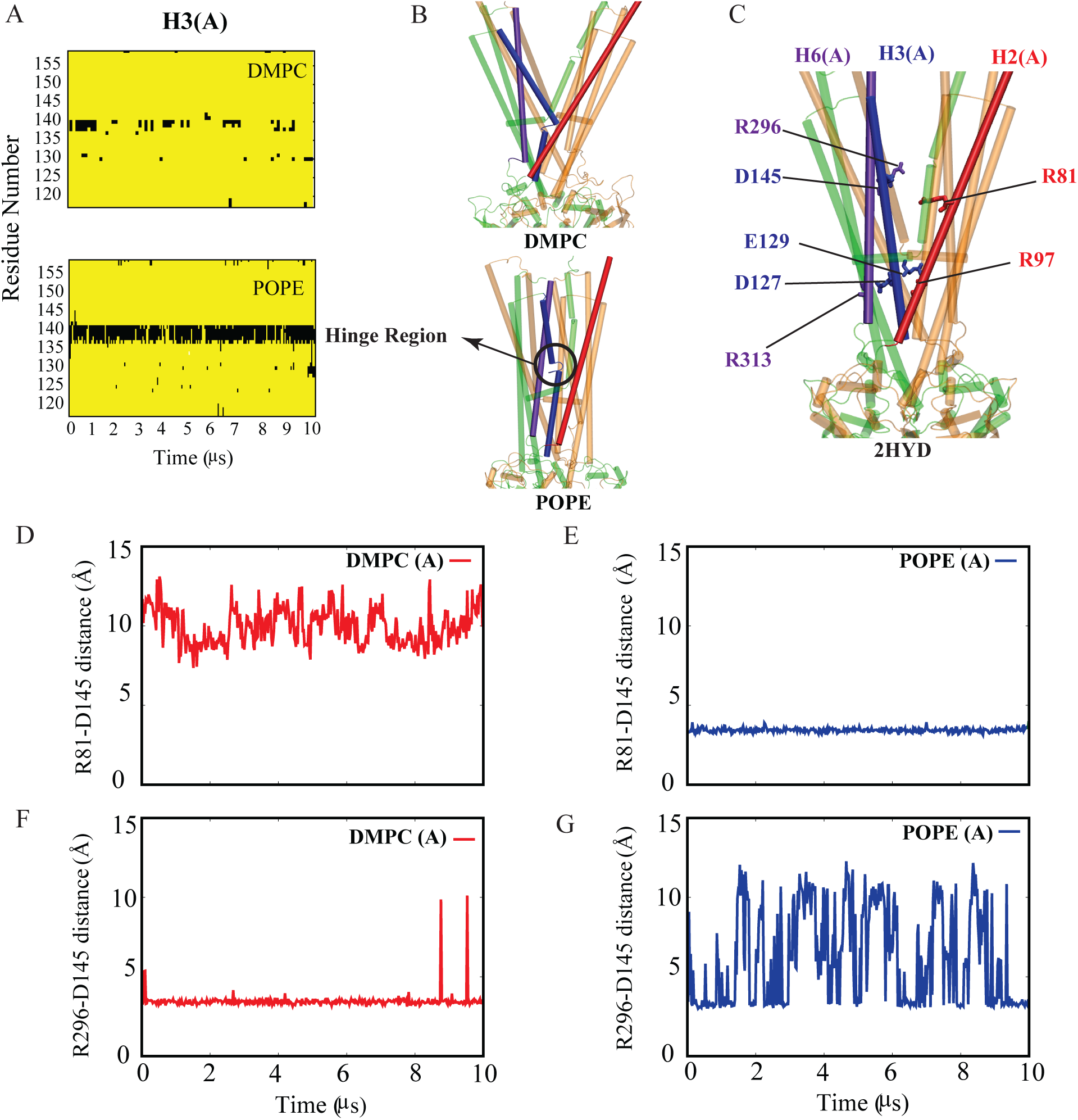
Absence of H3 Hinge Formation in DMPC Lipid Membrane: (A) secondary structure analysis displays the transformation of Helix 3 in monomer A of the protein in POPE, highlighting the transformation from helix (yellow color) to loop (black color). (B) presents the protein’s structure in the last frame within both DMPC and POPE lipid compositions, with the formation of the hinge region in POPE indicated by a black circle. (C) The initial structure of Sav1866 (2HYD) is depicted in this section, emphasizing the crucial role played by key residues in helices H2, H3, and H6 in the opening or closing of the periplasmic and cytoplasmic gates. (D, E) Salt bridge dynamics between residues R81 and D145 in DMPC and POPE, respectively, and in (F, G) between residues R296 and D145 in DMPC and POPE, respectively.

**FIG. 7.**
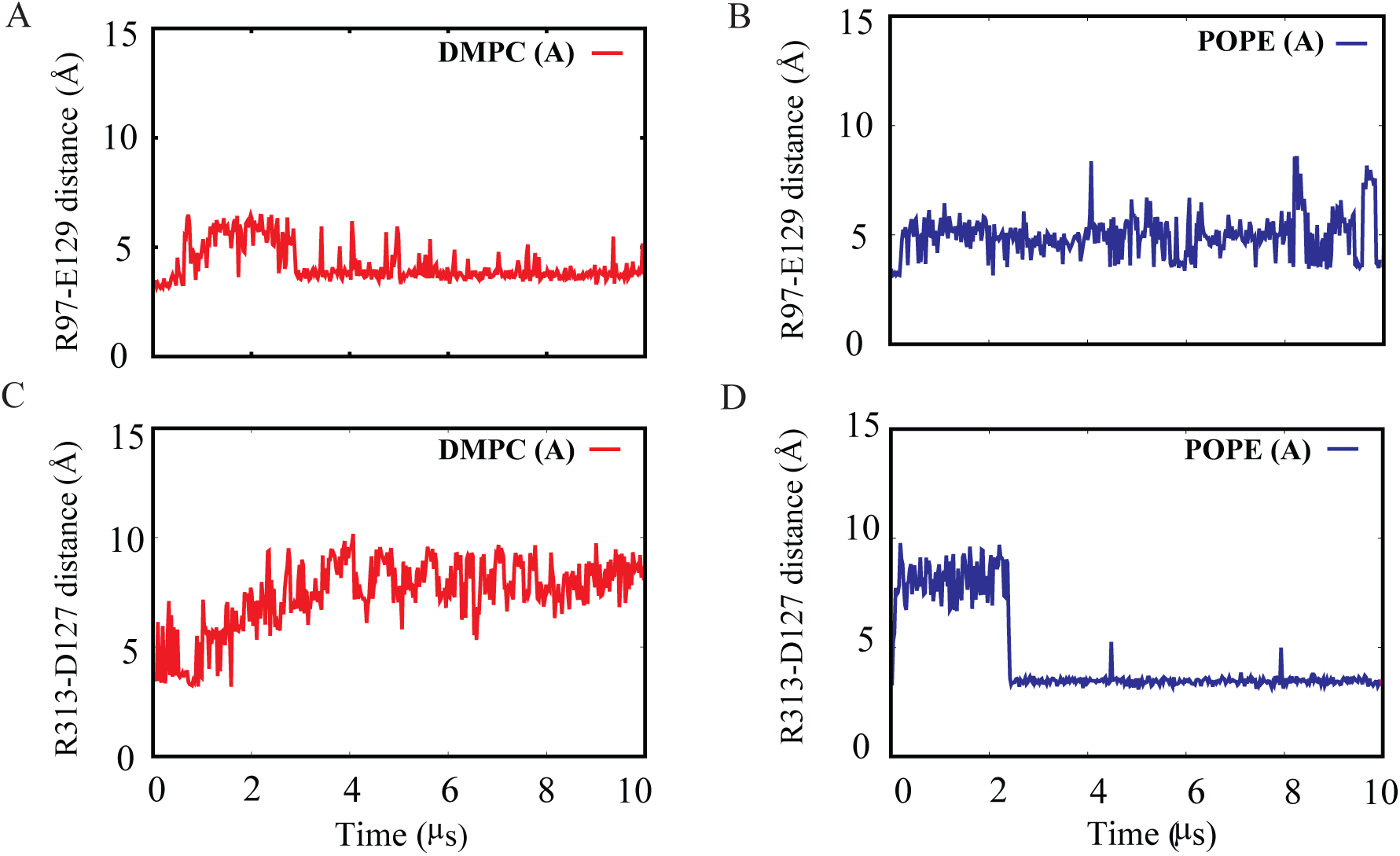
Illustration of the formation or breakage of Intrabundle and Interbundle salt bridges located below the hinge region of H3 in protomer A and adjacent to the cytoplasmic gate. (A, B) Interbundle salt bridge R97(H2)-E129(H3) in monomer A of the protein in DMPC and POPE simulations, respectively. (C, D) Intrabundle salt bridges between R313(H6) and D127(H3) in monomer A of DMPC and POPE simulations, respectively.

### Hydrogen Bond Interactions Influencing Gate Closure and Opening

In our earlier publication [1], it is hypothesized that the replacement of primary ammonium (PE) with quaternary ammonium (PC) in the lipid headgroups reduces their hydrogen bonding potential, which affects the conformational transitions of Sav1866, particularly in terms of the closure of the periplasmic gate. It is shown that within PE environments, the interactions between lipids and between lipids and the protein facilitate the closure of the transporter, while the presence of PC lipids at the periplasmic gate prevents its closure. These interactions of PE lipids trigger the formation of hydrogen bonds involving specific protein residues, such as D42(H1)-T276(H3) and K38(H1)-T279(H3).

In our current study, with extended simulations, we identified three stable hydrogen bond interactions within protein embedded into the POPE lipid that remained stable throughout the simulation, including previously described hydrogen bonds (D42-T276 and K38-T279), as well as an additional bond between M31 (H1) and Y286 (H6) on the periplasmic side, which was stable in at least one of the monomers (Figure. 8A, C). Importantly, these three hydrogen bonds are not formed in any of the monomers in the DMPC environment (Figure. 8B, D).

**FIG. 8.**
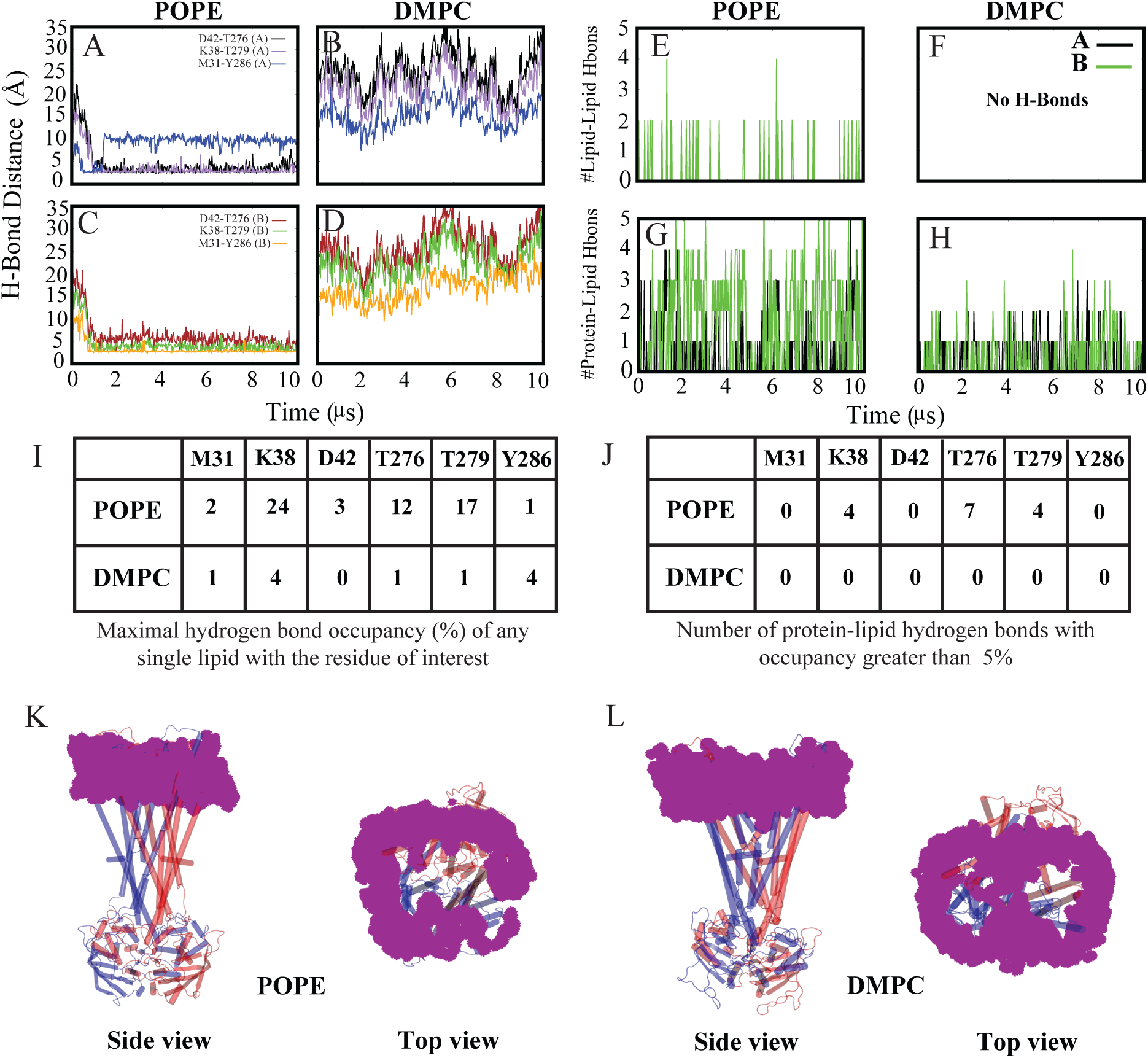
Lipid-mediated periplasmic gate closure. Hydrogen bond donor-acceptor distance time series for M31-Y286, K38-T279, and D42-T276 in monomer-A during POPE simulations (A) and DMPC (B) simulations. (C, D) Same as parts A and B for monomerB. Number of interlipid hydrogen bonds formed over time between lipids within 8Å of periplasmic residues M31, K38, D42, T276, T279, and Y286 are shown in POPE (E) and DMPC simulations (F). Number of lipid-protein hydrogen bonds formed over time between lipids and periplasmic residues M31, K38, D42, T276, T279, and Y286 are also shown in POPE (G) and DMPC (H) simulations. (I) displays the maximal hydrogen bond occupancy (in percentage) of any single lipid with residues of interest in both POPE and DMPC simulations. (J) Number of Protein-Lipid Hydrogen Bonds (Occupancy > 5%) are shown in both POPE and DMPC simulations. Outer leaflet lipid headgroup occupancy isosurfaces in and around the protein corresponding in POPE (K) and DMPC (L) simulations.

The lipid-protein hydrogen bond occupancy of POPE and DMPC lipids is shown in Figures.8 I. Residues M31, K38, and D42 from one TMD bundle (H1) and T276, T279, and Y286 from the opposite TMD bundle (H6) are identified as important contributors to the periplasmic closing, with their side chains engaging with PE lipid headgroups. In particular, these residues show minimal hydrogen bond formation with DMPC lipids. However, POPE lipids demonstrate the capacity to establish hydrogen bonds with residues situated at the periplasmic side of the transporter, leading to the closure of this gate.

While observing the behavior of outer leaflet lipids in proximity to the protein, a distinct contrast is noticed between POPE and DMPC simulations. In POPE simulation (Figure. 8K), all lipids exit the periplasmic gate, whereas in DMPC simulation (Figure. 8L), a substantial number of lipids remain in the vicinity of this gate. The presence of these lipids between the two transmembrane domain (TMD) bundles forming the periplasmic gate results in a blockage to the protein’s closure in a way that the primary ammonium in phosphatidylethanolamine (PE) promotes interactions between the two TMD bundles (Figure. 8E, G). In contrast, DMPC head groups lack the ability to form hydrogen bonds with each other, inhibiting the initiation of lipid-protein and protein-protein interactions (Figure. 8F, H).

When examining the cytoplasmic gate situated within the lower leaflet, our analysis of hydrogen bonds reveals significant distinctions between POPE and DMPC lipid environments. In POPE simulations (Figure. 9A), the interaction between lipid head groups is apparent, showing their ability to engage with each other. In contrast, in DMPC simulations (Figure. 9B), this interaction is missing, consistent with the expectation that DMPC head groups primarily serve as hydrogen bond acceptors. Furthermore, the hydrophobic tails of lipids, composed of long hydrocarbon chains, are not favorable for establishing hydrogen bonds. Consequently, the plots depicting lipid-lipid tail hydrogen bond interactions remain empty for both POPE and DMPC, as illustrated in Figure. 9E, F.

**FIG. 9.**
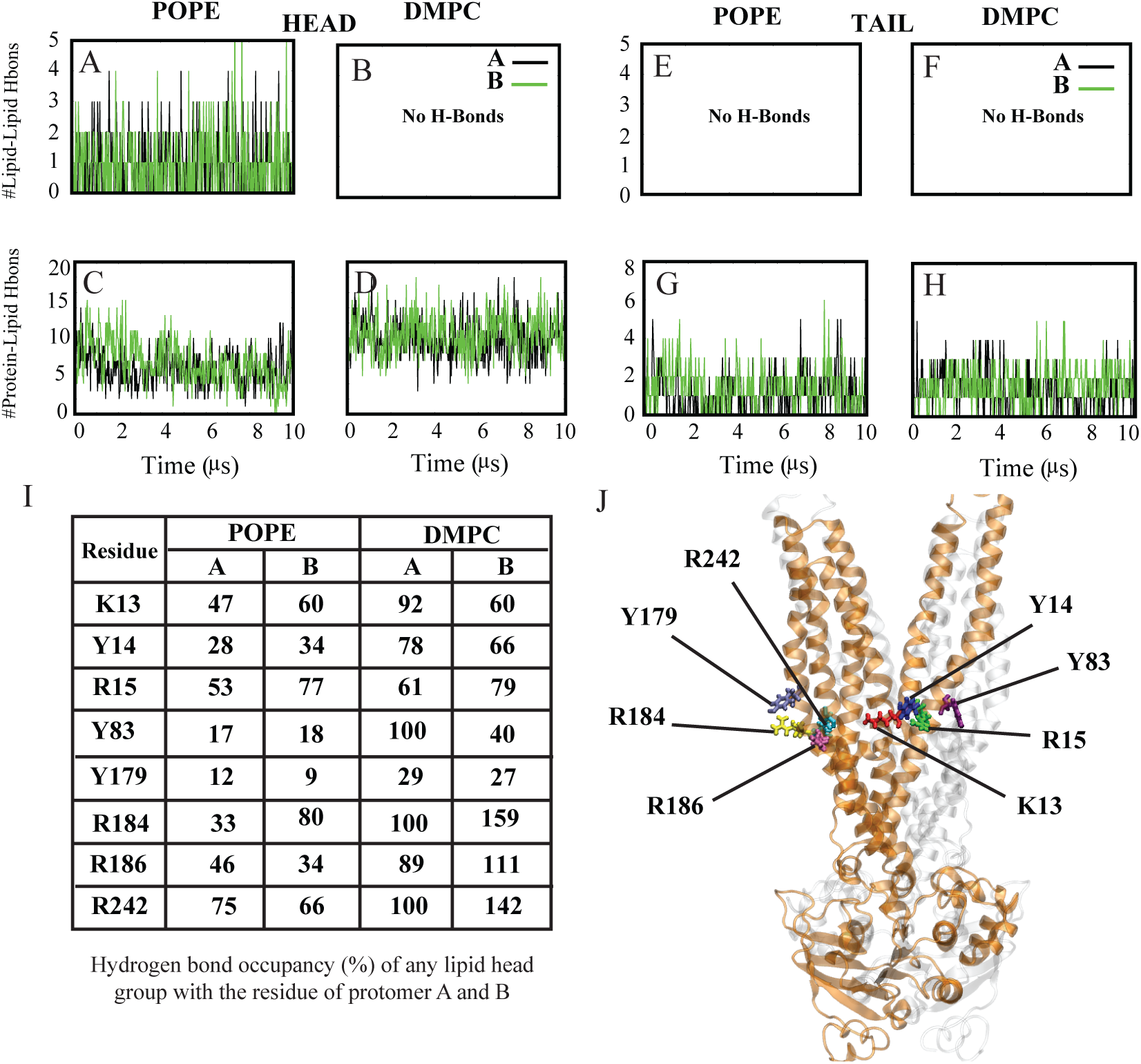
Lipid-Mediated cytoplasmic gate opening in DMPC. The increased formation of hydrogen bonds between the lipid head group and the protein within the cytoplasmic gate in DMPC is a critical factor triggering the opening of the cytoplasmic gate. Number of interlipid hydrogen bonds formed over time between lower leaflet lipids of head or tail within 8Å of protein in POPE (A, E) and DMPC simulations (B, F), respectively. Number of lipid-protein hydrogen bonds formed over time between lipid head or tails and protein are also shown in POPE (C, G) and DMPC (D, H) simulations. Number of hydrogen bonds in protomer A and B are shown with black and green colors, respectively. (I) Hydrogen bond occupancy (as a percentage) of any single lipid head with protomer A and B residues in both POPE and DMPC simulations. (J) Cartoon representation of the protein highlighting residues with higher interaction with lipids in monomer A.

In POPE, elevated lipid-lipid interactions between head groups correlate with a decrease in lipid-protein interactions (Figure. 9C), as acceptor and donor sites are involved in lipid-lipid interactions. In contrast, DMPC exhibits a lack of lipid-lipid interactions, allowing the head group oxygen (O) to act as an acceptor, leading to an increased interaction between lipid head groups and proteins (Figure. 9D). Furthermore, despite the longer tail in POPE compared to DMPC, protein-lipid tail hydrogen bonds exhibit similar behavior in both POPE (Figure. 9G) and DMPC (Figure. 9H).

Our investigation reveals specific residues (K13, Y14, R14, Y83, Y179, R184, R186, and R242) within the cytoplasmic gate that function as hydrogen bond donors, forming essential interactions with lipids and play a key role in the opening of the cytoplasmic gate. The occupancy of these hydrogen bonds between the protein and lipid headgroup in DMPC is higher than in POPE, particularly in monomer A (Figure. 9I).

As we move further down the pore, we observed the presence of two specific interbundle hydrogen bonds (ASP319-HSD103 and ARG97-VAL124) between H2 and H3 or H6 with significantly high occupancy in the protein embedded in POPE, which were not formed in the DMPC environment (Figure. 10A). Upon examining the interactions between lipid headgroups and the protein, as well as the interactions of lipid headgroups within an 8Å of these four residues, we realized that in the DMPC environment, there are no interactions between lipid molecules (Figure. 10D), or lipid and proteins (Figure.10F), whereas in the POPE environment, lipid headgroups engaged in hydrogen bonding with both lipids (Figure. 10C) and protein residues (Figure. 10E). The longer tails in POPE bring the lipid heads closer to protein residues, facilitating the formation of hydrogen bonds between them. These interactions effectively brought the residues close together, leading to the formation of the hydrogen bonds in POPE. The absence of such interactions in DMPC contributed to the opening of the pore.

**FIG. 10.**
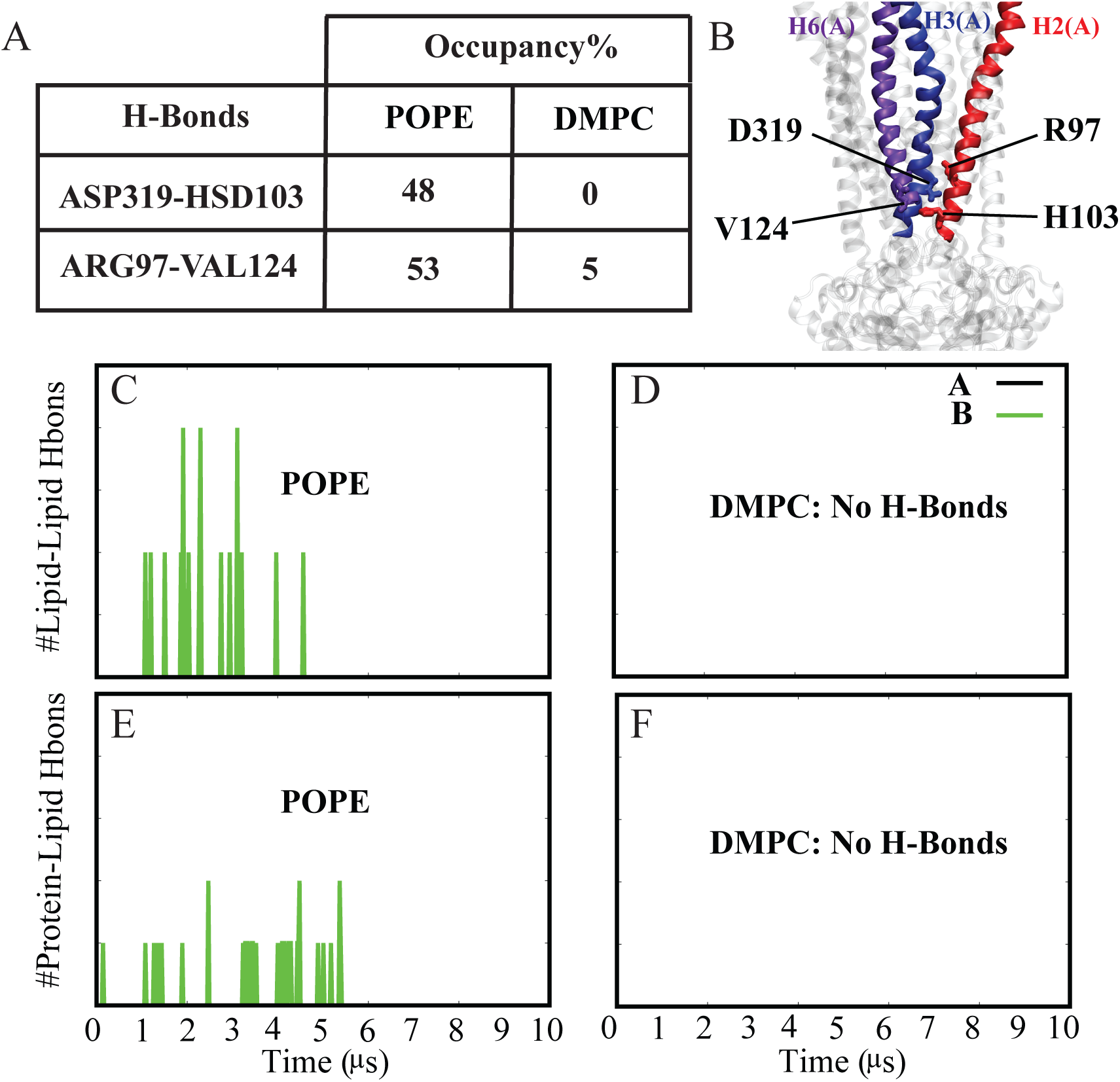
Formation of h-bonds between H2, H3, H6 in POPE. Two interbundle hydrogen bonds (ASP319(H3)-HSD103(H2) and ARG97(H2)-VAL124(H6)), which were not formed in the DMPC environment, exhibit significantly high occupancy in the protein embedded in POPE. (A) The table displays the occupancy of hydrogen bonds in POPE and DMPC environments. (B) Showing the location and helix number of residues responsible for the identified hydrogen bonds. (C) Depicts the number of interlipid hydrogen bonds as a function of time formed between lipids positioned within 8Å of the cytoplasmic residues D319, H103, R97, and V124 in POPE. (D) Illustrates the same for DMPC simulations. (E) Presents the number of lipid-protein hydrogen bonds as a function of time formed between lipids and cytoplasmic residues D319, H103, R97, and V124 in POPE. (F) Shows the same for DMPC simulations. Green color indicates hydrogen bonds in protomer A, while black color represents hydrogen bonds in protomer B in all four plots.

### Flippase activity of Sav1866 in DMPC simulation

Flippases facilitate the bidirectional translocation of lipids between the two leaflets of a lipid bilayer. This activity has been observed in both PO (e.g., POPE2) and non-PO (e.g., DOPE2 and DOPE3) simulations in our previous study on Sav1866 [1]. In these lipid systems, the protein catalyzes the movement of lipids between leaflets on relatively short time scales. This activity is carried out by the charged side chains located in or near the translocation chamber, which exhibited interactions with lipid headgroups. These interactions were found to inhibit the closure of the periplasmic gate, a phenomenon observed in non-POPE lipid environments, with an important caveat that the periplasmic gate closure was not always blocked (POPE), as lipids could align themselves along the membrane’s normal axis. In our new study, we have observed flippase activity within the DMPC lipid system (non-PO). This new finding aligns with our previous observations that the flippase activity in non-PO systems prevents the periplasmic closure. Our investigations demonstrate that the flippase activity involving DMPC occurs approximately 9 *µ*s after exposure, and it is coordinated through interactions with positively charged residues, specifically R295, R296, and R81, and the negatively charged phosphate groups present in the DMPC headgroup. In comparison to the POPE simulation, the analysis indicates that the flipped lipid exhibits reduced interaction with periplasmic gate residues (K38 and T279) in the DMPC lipid environment. At the same time, the flipped lipid forms new hydrogen bonds with positively charged residues in the translocation chamber (R295, R81, and R296) (Figure. 11B).

**FIG. 11.**
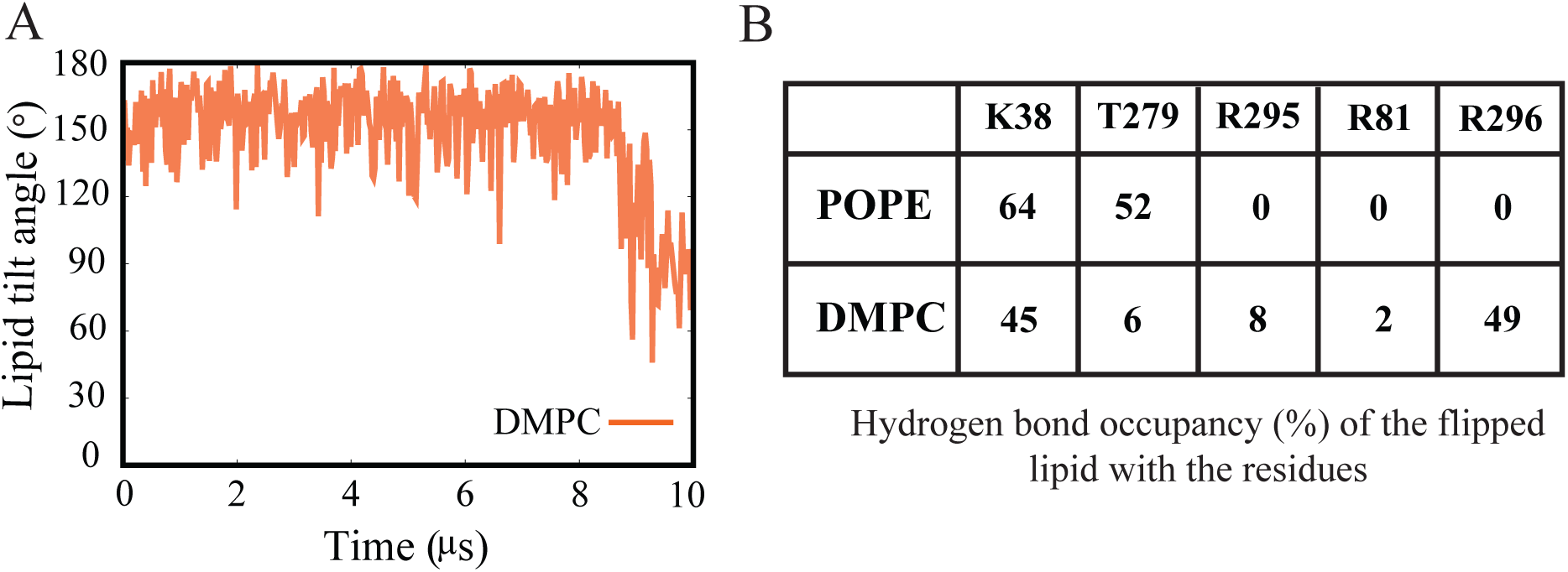
Flippase activity of Sav1866 in DMPC simulation. (A) Tilt angle of the flipped lipid in DMPC as a function of time. (B) Hydrogen bond occupancy % of periplasmic residue (K38 and T279) and residues located in the translocation chamber (R295, R81, and R296) with flipped lipid.

In the previous study [1], it was suggested that E288 acted as the primary driving force to attract PE lipids into the translocation chamber of the transporter. This hypothesis was based on the idea that E288, positioned slightly above the membrane’s center, had a higher chance of interacting with the headgroups of the outer leaflet lipids headgroup and positively charged primary ammonium groups of PE lipids, which can form hydrogen bonds with E288.

In contrast to the findings of the previous study, where it was proposed that PC lipids, lacking primary ammonium groups in their headgroups, were unable to interact with E288 and, as a result, did not enter the chamber from the center nor undergo flipping, our latest research has uncovered a different scenario. Our observations indicate that PC lipids can enter the chamber. However, this process occurs with a notable time delay, taking approximately 9 *µ*s (Figure. 11A). This significant delay challenges the earlier assumption that PC lipids remained excluded from the translocation chamber due to their lack of interaction with E288. The extended timescale for their entry into the chamber suggests a more complex and time-dependent interaction mechanism.

### NBD-NBD interface

After removing ATP from the NBDs, the expected behavior was that the two NBDs would completely dissociate due to ATP hydrolysis and ADP release. To investigate this, we calculated the distance between the center of masses of the two NBDs. The results (Figure. 12A) indicated an initial increase in the distance between the two NBDs in both DMPC and POPE during the first 5 *µ*s of the simulation. However, after 5 *µ*s, they exhibited different behaviors, with one NBD distance decreasing (POPE) while the other increased (DMPC). Additionally, we measured the *γ* angle (Figure. 12B), which measures the angle between the two NBDs and indicates the twisting of the NBDs relative to each other (Figure. 2). This angle displayed a similar pattern in both lipid environments.

**FIG. 12.**
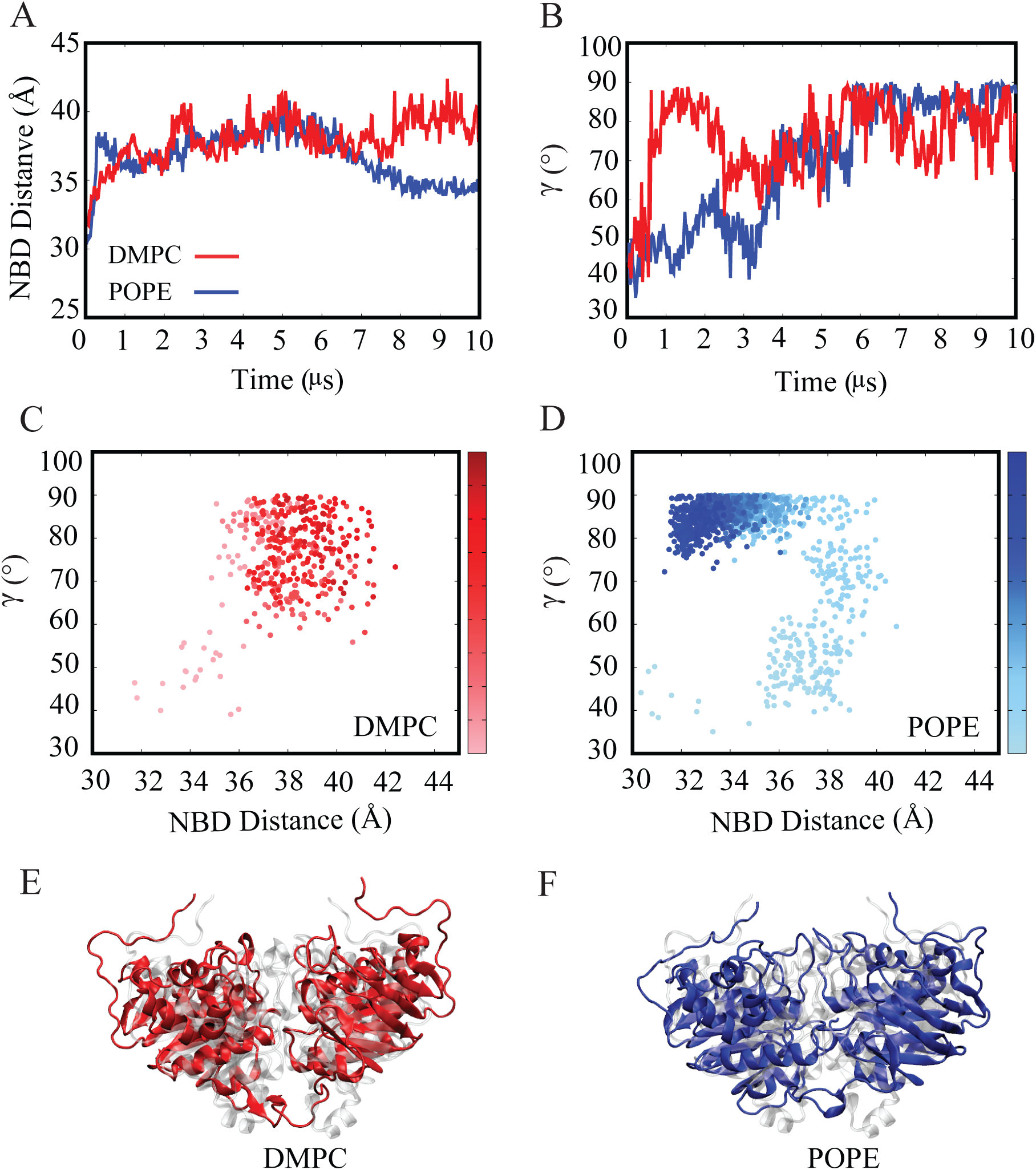
(A) Presents the time series of the NBD (Nucleotide-Binding Domain) distance, and (B) depicts the *γ* angle (twisting angle) in the presence of both POPE and DMPC. The data for POPE is represented in blue, whereas DMPC data is displayed in red.(C) and (D) display the outcome of projecting the gamma and NBD distance in relation to each other. (E & F) Cartoon representation of the protein’s NBD domain embedded in DMPC and POPE, respectively. The molecular images are compared with the first frame of this domain represented in gray.

In the POPE environment, the NBD distance initially increased (Figure. 12A) and then decreased, while the *γ* angle started at 45*^◦^* and increased to 90*^◦^*, remaining constant (Figure. 12B). This suggests that NBD dimerization occurs in a specific orientation of 90*^◦^* in POPE. In contrast, in the DMPC environment, the NBD distance continued to increase (Figure. 12A), and the twisting angle fluctuated around 90*^◦^* (Figure. 12B). The projection of NBD distance and *γ* angle (Figure. 12C, D) clearly illustrates that the protein’s behavior undergoes a distinct transformation in response to different lipid environments. In the POPE lipid environment (Figure. 12D), the protein initially transitions from the outward-facing (OF) state, characterized by a distance of 33 Å and an angle of 45*^◦^*, to the occluded inward-facing (IF-occ) state, by a distance of 35 Å and a *γ* angle of 90*^◦^*. There are also intermediate states identified in between these extreme conformations. Intermediate states were observed, characterized by distances of 37 Å with a *γ* angle of 50*^◦^*, and another at 39 Å with a *γ* angle of 75*^◦^*.

In the crystal structure of the outward-facing (OF) state of Sav1866, a pattern of positively and negatively charged residues is observed in the NBD-NBD interface (Figure. S6).

These charged residues exhibit distinct interactions when the NBDs are positioned in specific orientations. Therefore, we propose that alterations in the distance between these charged residues have a direct impact on the *γ* angle, which serves as a key determinant of the required orientation. Once the NBDs reach this specific orientation, they form the interactions, facilitating a new dimerization process that results in a reduction in NBD distance. Our hypothesis aligns with the hypothesis presented in Moradi et al.’s work [39], which suggests that charged residues within the NBD-NBD interface play an important role in facilitating the OF to IF transition in the multidrug transporters.

In Figure. 13A, we have visualized the charged residues within one monomer of the NBD-NBD interface, revealing their specific orientation. Positively and negatively charged residues are arranged side by side in a direction spanning from 2 o’clock to 8 o’clock. In this arrangement, the other monomer initially slides into position and subsequently locks into place by interacting with oppositely charged residues (Figure. 13C).

**FIG. 13.**
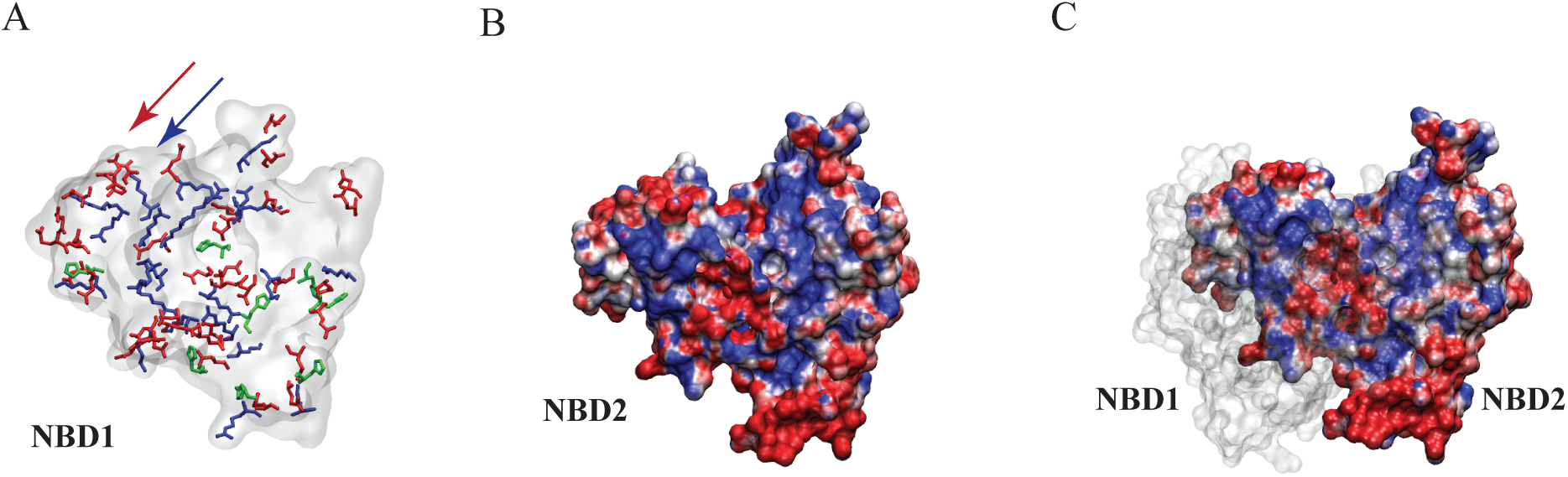
NBD-NBD Interface. (A) indicates the distribution of positively and negatively charged residues at the dimeric interface of one monomer, highlighted in blue and red, respectively. Histidines are also shown in green. (B) illustrates the surface charge distribution of one NBD on the NBD-NBD interface, as captured from a snapshot of the Sav1866 structure in the OF conformation. Blue and red indicate positive and negative charges, respectively. (C) demonstrates the NBD-NBD interaction.

Figure. S6B indicates the NBD dimer structure at the last frame of a 10 *µ*s simulation in the POPE environment. Here, it becomes apparent that the charged residues can interact with one another in an altered orientation compared to the initial configuration. However, in the case of DMPC (Figure. S6C), the orientation of these charged residues results in repulsion. This is evident from the presence of blue spots in the figure, which indicate an accumulation of positively charged residues contributing to this repulsive effect. Consequently, the NBD distance increases and the *γ* angle continues to fluctuate as the NBDs seek the optimal orientation for interaction. Therefore, it becomes evident that the charged residues located within the NBD-NBD interface have a significant influence on the NBD twist and separation. This influence can likely be attributed to the disruption of the NBD twist mechanism, which is primarily governed by the electrostatic interactions occurring within the NBD-NBD interface.

### Sav1866 Remains in the Occluded Inward-Facing State in Extended POPE Simulations

In order to gain deeper insights into the protein’s conformational changes, we extended the simulation to 30 *µ*s in the POPE environment.

During this extended period, the protein exhibited remarkable stability, as evidenced by the consistently low RMSD values (Figure. S10), along with the constant *α* and *β* angles (Figure. S11). Furthermore, a comprehensive analysis, including water density maps, lipid density, and secondary structure, was conducted, all of which contributed to a detailed understanding of the protein’s behavior.

Among these analyses, a significant finding was the reduction of the NBD distance to approximately 32 Å(Figure. 14A). This reduction in distance suggests that the NBDs reached and maintained a specific orientation favorable for stable NBD-NBD interactions (Figure. 14B). This observation provides valuable insights into the underlying mechanisms of the protein’s stability and function. Our research revealed the formation of two salt bridges (GLU570-ARG485 and GLU513-ARG535) within the NBD-NBD interface that are stable for at least 5 *µ*s of the simulation. These findings shed light on the structural and electrostatic factors contributing to the protein’s stability and functionality in the occluded inward-facing state (IF-occ).

**FIG. 14.**
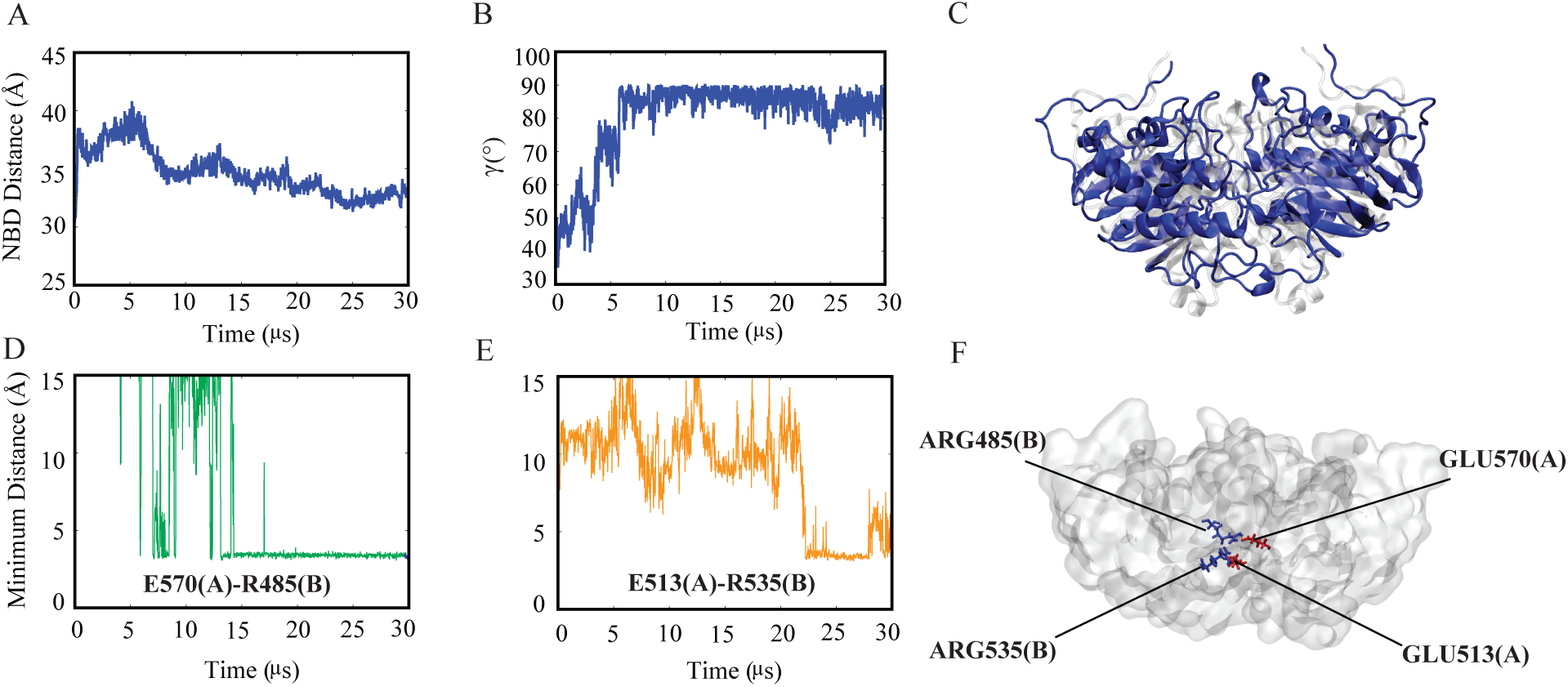
(A) shows the distance between the center of mass of two NBDs of the protein embedded into a POPE lipid composition at the 30 *µ*s of the simulation. (B) illustrates the twist or gamma angle of the NBDs at the same 30 *µ*s time point in the simulation. (C) Cartoon representation of the protein’s NBDs in the last frame of the 30*µ*s simulation, embedded in POPE (blue), compared to the first frame of this domain, depicted in gray. (D & E) Salt Bridge Interactions between two NBDs in POPE Simulation during 30*µ*s: Negatively charged residues GLU570 and GLU513 from monomer A establish stable salt bridge interactions with positively charged ARG485 and ARG535 from monomer B, respectively. These interactions form the NBD-NBD interface, contributing to the overall stability of the system. (F) key residues responsible for the stable NBD-NBD interface through the salt bridge interactions are highlighted.

We investigated the distribution of charges on the NBD-NBD interface in the POPE environment (Figure. 15). We observed positive charges (blue), negative charges (red), and histidine residues (green) at various times during the 30 *µ*s simulation. These charges demonstrated dynamic interactions that influenced NBD dissociation and reassociation, including repulsion and attraction, respectively. Initially, the NBD-NBD interface had a distinct configuration of positive and negative charges (Figure. 15A). However, as the simulation progressed, the charges reorganized, possibly as a result of repulsive interactions between similarly charged residues (Figure. 15B). Interestingly, the distribution of charges appeared to influence the orientation and relative position of the NBDs (Figure. 15C, D, and E). This dynamic interaction influenced the closure of the cytoplasmic gate. Furthermore, the observed twisting angle and changes in the surface charge distribution suggested a complex interaction between charge distribution and NBD reassociation. At certain orientations, the charges facilitated NBD dimerization, bringing the NBDs closer together. Repulsive interactions between charged residues, on the other hand, could have contributed to NBD dissociation. The NBDs continuously changed their relative orientation during the simulation in an attempt to find the ideal configuration, possibly to minimize repulsive interactions and facilitate reassociation.

**FIG. 15.**
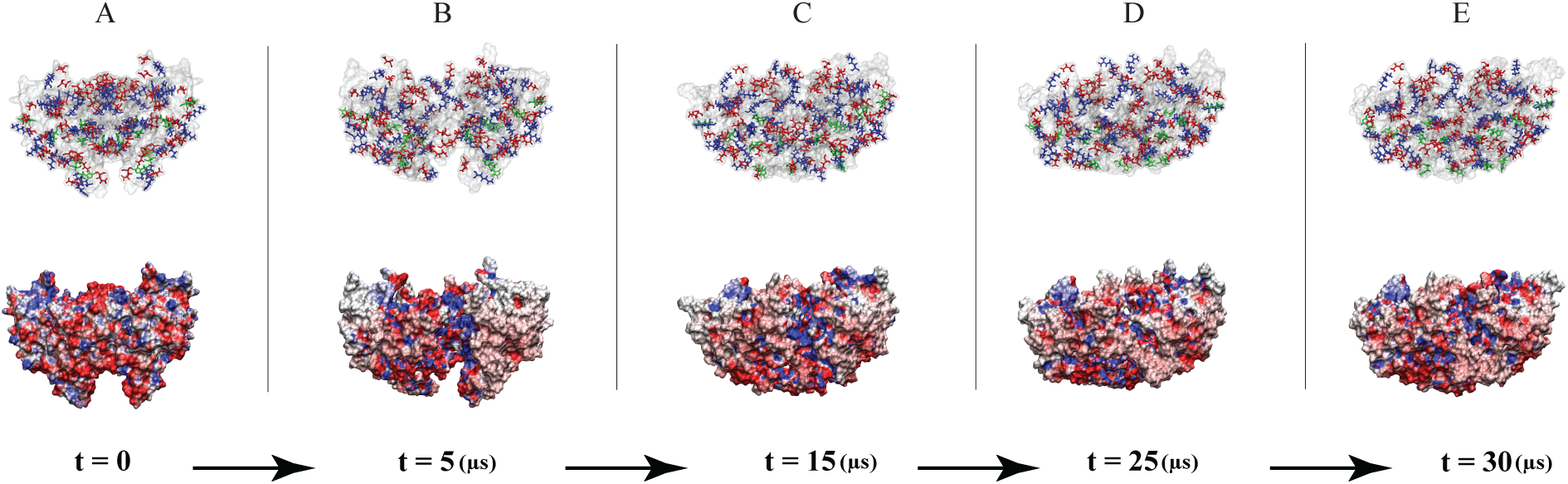
NBD-NBD Interface during a 30 *µ*s simulation of POPE. The top arrow illustrates the distribution of positive (blue), negative (red), and histidine (green) charges on the dimeric interface at various time points (0, 5, 15, 25, and 30 *µ*s). The bottom arrow indicates the surface charge distribution for the NBD interface, with blue and red representing positive and negative charges, respectively.

## DISCUSSION

In this comprehensive study, we employed microsecond-level all-atom equilibrium molecular dynamics (MD) simulations to investigate the structural dynamics of the bacterial ABC exporter Sav1866 in different lipid environments, specifically POPE and DMPC. Our findings reveal crucial insights into the lipid-dependent conformational dynamics of ABC transporters, emphasizing the specificity of lipid interactions in shaping the function and mechanism of these transporters. One of the most remarkable findings in our study is the distinct behavior of Sav1866 when embedded in the DMPC lipid composition. While in POPE, Sav1866 undergoes a transition from the outward-facing (OF) to the inward-facing occluded (IF-occ) state, we propose an unconventional transition in DMPC, deviating from the traditional Alternating Access Mechanism. In DMPC, Sav1866 exhibits a unique “channel-like” behavior, with both cytoplasmic and periplasmic gates open throughout the 10 *µ*s simulation. The presence of three periplasmic interbundle hydrogen bonds (D42(H1)-T276(H3), K38(H1)-T279(H3), and M31(H1)-Y286(H6)) and lipid-lipid interactions in POPE, absent in DMPC, and higher lipid-protein interaction in POPE, illustrates the impact of lipid interactions on protein conformations. These interactions in POPE facilitate the closure of the periplasmic gate by bringing the protein closer. In contrast, with respect to the cytoplasmic gate of the protein, DMPC exhibits stronger interactions with the protein compared to POPE. These interactions exert an outward force on the protein, resulting in the opening of the cytoplasmic gate. Our investigation identifies specific residues (K13, Y14, R14, Y83, Y179, R184, R186, and R242) within the cytoplasmic gate that serve as hydrogen bond donors. These residues establish crucial interactions with lipids, playing a crucial role in the opening of the cytoplasmic gate. The occupancy of these hydrogen bonds between the protein and lipid headgroup in DMPC is significantly higher than in POPE, particularly in monomer A. Furthermore, as we explore the pore region, the absence of two critical hydrogen bonds (ASP319(H3)-HSD103(H2) and ARG97(H2)-VAL124(H6)) in DMPC, compared to POPE, significantly influences the opening of the cytoplasmic gate. The longer tail in POPE allows for closer proximity to these residues, promoting hydrogen bond formation. Interestingly, our study also reveals partial flippase activity in DMPC, where lipids from the outer leaflet of the membrane enter the substrate translocation chamber, attracted by positively charged residues. This activity becomes evident after 9 *µ*s of simulation. In addition, the NBD-NBD interface displays a distinctive arrangement of positively and negatively charged residues. When precisely oriented, these residues form bonds with each other. Notably, the simulation in POPE for 30 *µ*s reveals the formation of salt bridges (GLU570-ARG485 and GLU513-ARG535) between the two NBDs. This interaction plays a pivotal role in reducing the NBD distance. Our research highlights the complex relationship between lipid content and Sav1866’s conformational dynamics. Lipid interactions have a crucial role in regulating the activity of ABC transporters, as evidenced by lipid-specific characteristics such as the channel-like behavior in DMPC and the transition to IF-occ in POPE. This work opens the door for more research into the lipid selectivity of membrane proteins by providing insightful information on the complex relationship between lipid environments and the conformational dynamics of ABC transporters.

## EXPERIMENTAL PROCEDURES

It is computationally challenging to characterize membrane transporter structural transitions without sacrificing a detailed chemical description of these systems and the environment in which they operate, primarily since such processes demand prohibitively long times. We used all-atom MD to investigate the OF-IF conformational transitions of Sav1866 transporters embedded in different lipids in the apo state. Our simulations were based on X-ray structure of the Sav1866 transporter (PDB entry:2HYD) [11] The simulation input generator CHARMM GUI [13, 23] was employed to build each lipid-protein system. Each simulation system comprises one protein, *^∼^*= 360 lipids, 0.15 M NaCl, and *^∼^*= 45000 TIP3P water molecules [14]. The overall size of the system was *^∼^*= 134 × 142 × 174 Å ^3^. To describe all molecules, we employed the CHARMM36 all-atom additive force field parameters [16–18, 44]. NAMD 2.10 [15, 43] was used for preliminary MD simulations before Anton 2 production runs. A Langevin integrator was used to simulate each system with periodic boundary conditions at 310 K temperature, 0.5/ps collision frequency, and a time step of 2 fs. The pressure was maintained at 1 atm using the Nos’e-Hoover Langevin piston method [19, 20]. The smoothed cutoff distance for non-bonded interactions was set to 10-12 Å, and the long-range electrostatic interactions were calculated using the particle mesh Ewald (PME) method [21]. Prior to equilibration, each system was energy minimized for 10 000 steps using conjugate gradient algorithm [22] and further relaxed using a multistep restraining procedure (CHARMM-GUI’s default procedure for membrane proteins) [23] for *∼*1 ns. The initial relaxation was performed in NVT ensemble, followed by a 5 ns equilibrium simulation in the NPT ensemble. Structures have been obtained from each 5 ns preliminary simulation for the 10 *µs* and 30 *µs* production runs on Anton 2 for DMPC and POPE, respectively. Every production run was conducted with a 2.5 fs time step. The pressure was maintained semi-isotropically at 1 atm using the MTK barostat, and the temperature was controlled at 310 K using the Nosé-Hoover thermostat [19, 20]. The fast fourier transform (FFT) method [24], implemented on Anton 2, was used to compute the long-range electrostatic interactions. The configurations were collected every 2.4 ns. To analyze, create molecular snapshots, and visualize the data, visual molecular dynamics (VMD) was utilized [25]. We utilized VMD and its numerous plug-ins to calculate hydrogen bonds, salt bridges, secondary structure, membrane parameters (such as thickness, interdigitation, deuterium order parameter (SCD), and tilt angle), and water and lipid densities [25, 27, 40]. TM helices H1, H2, H3, H4, H5, and H6 are made up of residues 1–44, 52–106, 117–159, 161–216, 218–272, and 277–319 in each monomer. The residues 321–578 of each monomer compose each NBD. Bundle 1 is made up of monomerA’s residues 1–107 (TM helices H1 and H2) and monomerB’s residues 116–320 (TM helices H3–H6). Conversely, bundle 2 comprises residues 1–107 from monomerB and 116–320 from monomerA. Interbundle angle *β* and *α* was calculated as the angle between the third principal axes of the two TMD bundles in OF and IF conformation, respectively. To estimate the likelihood of finding water and lipid molecules near and inside the protein, we used density isosurface calculations. We examined cross sections of these density isosurfaces for water to determine its accessibility within and around the protein in different lipid environments. These cross sections were taken along the yz plane, with a 10 Å slice considered along the x-axis. When generating lipid density profiles, we only considered lipids in the outer leaflet within 10 Å of the protein. The cutoff distance and angles evaluated in the hydrogen bond study were 3.5 Å and 30*^◦^*, respectively. The smallest distance between donor and acceptor atoms was used to compute salt bridge distances. The PRODY software was used to do principal component analysis (PCA) [28]. For the PCA calculations, only C*α* atoms were used. The distances between the two NBDs were estimated using the centers of mass of their C*α* atoms. For lipid-related analyses, including Area per lipid (APL), membrane thickness, lipid tilt angle, and SCD, we employed the MEMBPLUGIN within VMD [40]. This tool facilitated comprehensive investigations into various lipid properties, enhancing our understanding of the membrane structure and dynamics in the context of our study. The lipid tilt angle was measured as the change of vector connecting the phosphorus atom in the headgroup to the terminal carbon of the acyl chains.

## Data Availability

https://github.com/bslgroup/sav1866

## Supporting Information

This article contains Supporting Information.

## Acknowledgements

This research is supported by the National Science Foundation (NSF) CAREER award (CHE 1945465), the National Institutes of Health grant R15GM139140, R35GM147423, and the Arkansas Biosciences Institute. This work also used the Extreme Science and Engineering Discovery Environment (XSEDE), which is supported by National Science Foundation Grant ACI-1548562. This work used XSEDE resources Comet and Stampede through allocation MCB150129. Anton 2 computer time was provided by the Pittsburgh Supercomputing Center (PSC) through Grant R01GM116961 from the National Institutes of Health. The Anton 2 machine at PSC was generously made available by D.E. Shaw Research.

## Notes

### Competing Interest Statement

The authors have declared no competing interest.

### Summary of Updates

The name of the first author has been corrected.

