## Supporting Information for "Lipid-dependent conformational dynamics of bacterial ATP-binding cassette transporter Sav1866"

### Supporting Figures

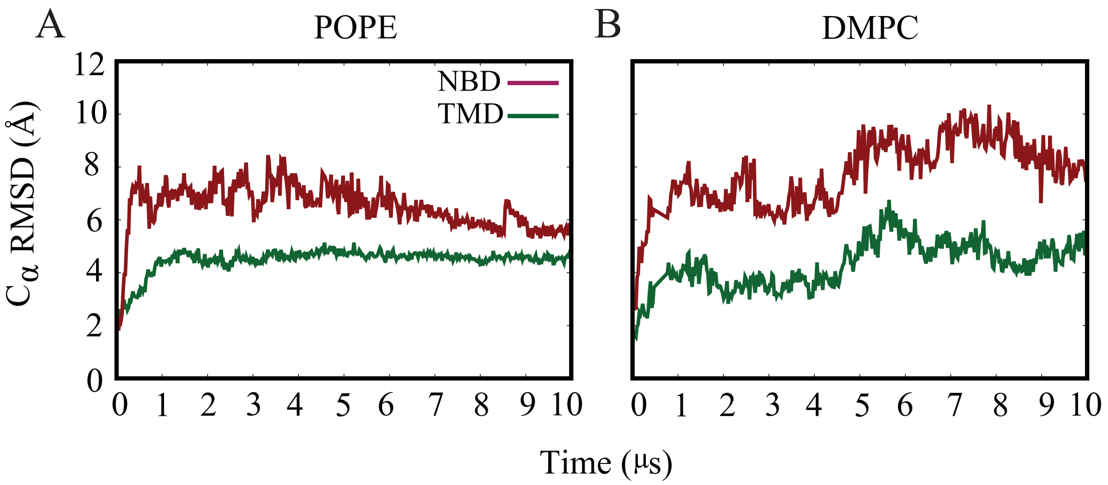


**Figure S1**: **C$\alpha$ RMSD for individual protomers.** The internal C-$\alpha$ RMSD calculated for NBD and TMD relative to their initial crystal structure over the 10 $\mu$s simulation is plotted for both POPE (A) and DMPC (B) systems. NBD is colored dark brown, and TMD is colored dark green. The TMD is more stable overall than the NBD.

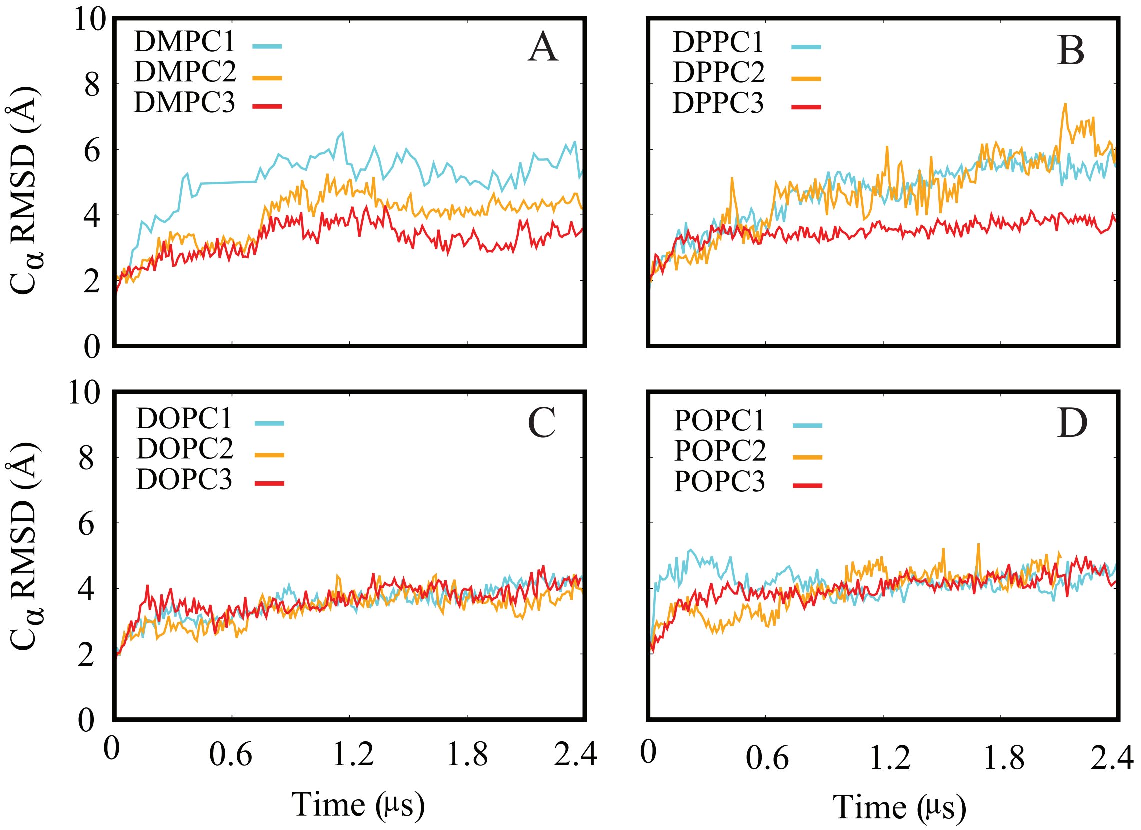


**Figure S2**: **C$\alpha$ RMSD in various lipid PC compositions.** Time series of root mean square deviation (RMSD) of C$\alpha$ atoms of the protein with respect to the initial model. The RMSD time series associated with the apo protein in different lipid environments, including DMPC (A), DPPC (B), DOPC (C), and POPC (D), are presented in panels A to D, respectively. Each color represents a repeated system, providing insights into the structural dynamics of the protein under various lipid compositions over 2.4$\mu$s of the simulation.

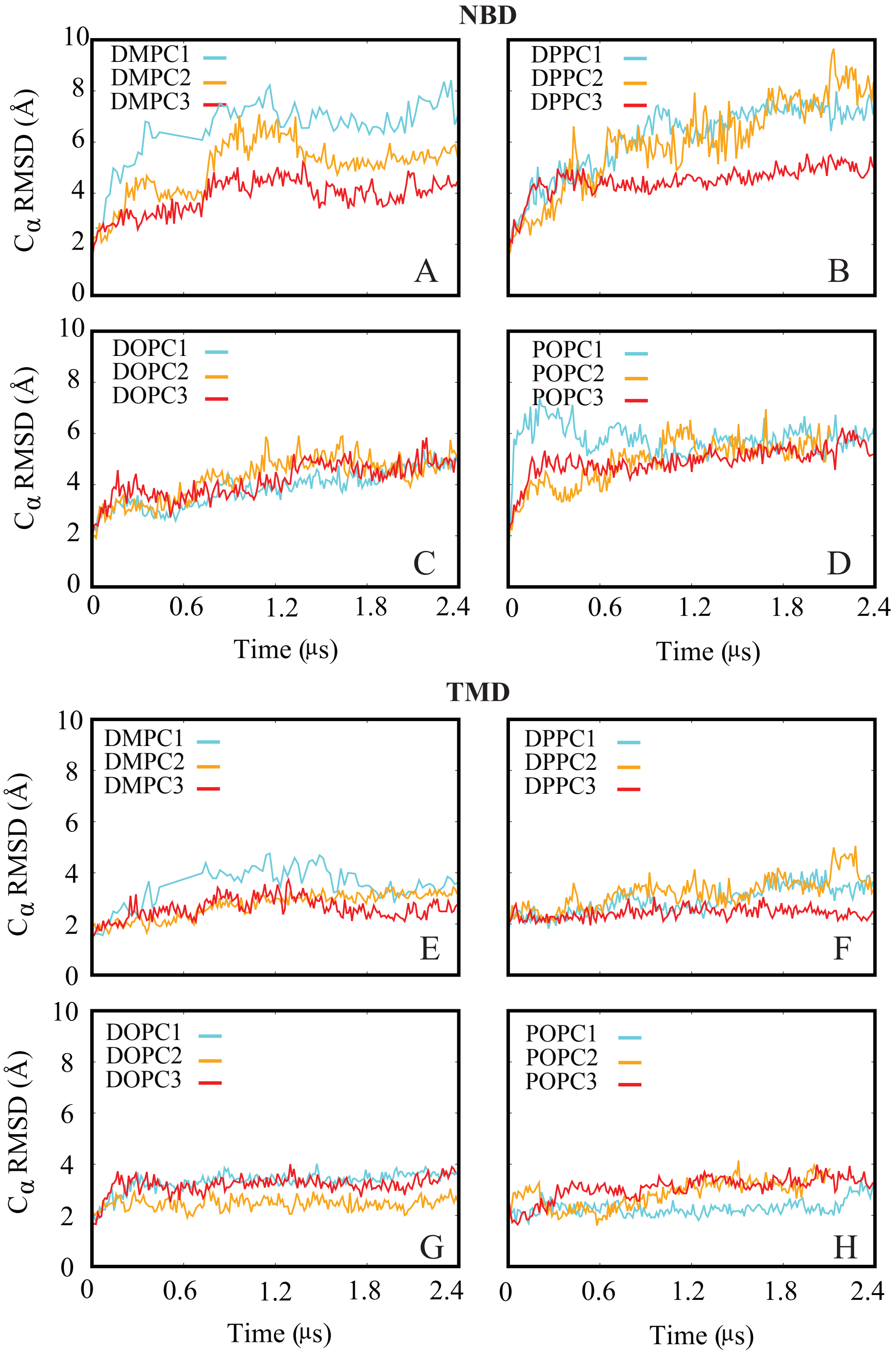


**Figure S3**: **Internal C$\alpha$ RMSD of NBD and TMD in various PC lipid compositions.**C-$\alpha$ RMSD for individual nucleotide-binding domains (NBD) and transmembrane domains (TMD) in various phosphatidylcholine (PC) lipid compositions. The internal C$\alpha$ RMSD, calculated for NBDs relative to their initial crystal structure over the 2.4 $\mu$s simulation, is depicted for DMPC (A), DPPC (B), DOPC (C), and POPC (D) systems. Subsequently, panels (E), (F), (G), and (H) illustrate the internal C$\alpha$ RMSD calculated for TMDs in DMPC, DPPC, DOPC, and POPC systems, respectively. This comprehensive analysis provides insights into the dynamic behavior of both NBD and TMD regions across different PC lipid compositions.

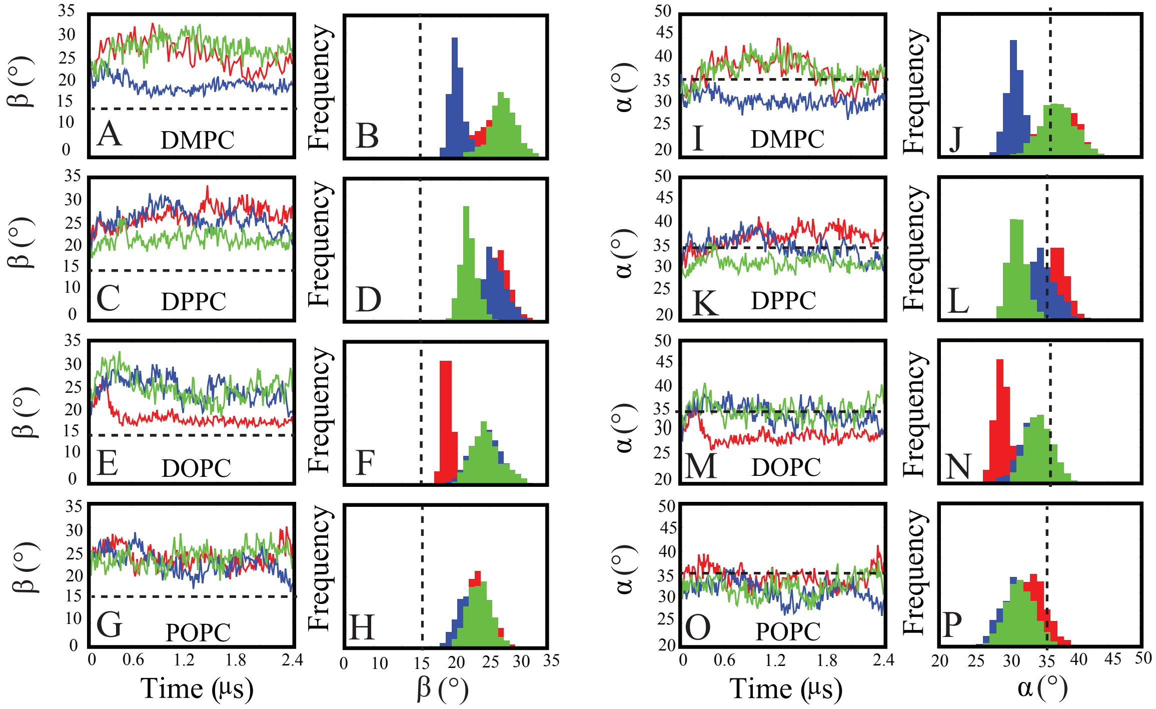


**Figure S4: Time series and frequency distributions of $\beta$ angles (A-H) and $\alpha$ angle (I-P) for the PC systems over a 2.4 $\mu$s simulation period in each repeat.** The protein exhibits consistent opening from the periplasmic gate, as indicated by the behavior of the $\beta$ angle in all PC simulations (A-H). However, the cytoplasmic gate, characterized by the $\alpha$ angle, is open in the two repeats of DMPC simulation, as observed in the frequency plot. In contrast, the cytoplasmic gate ($\alpha$) tends to be closed in the majority of repeats for DPPC, DOPC, and POPC simulations.

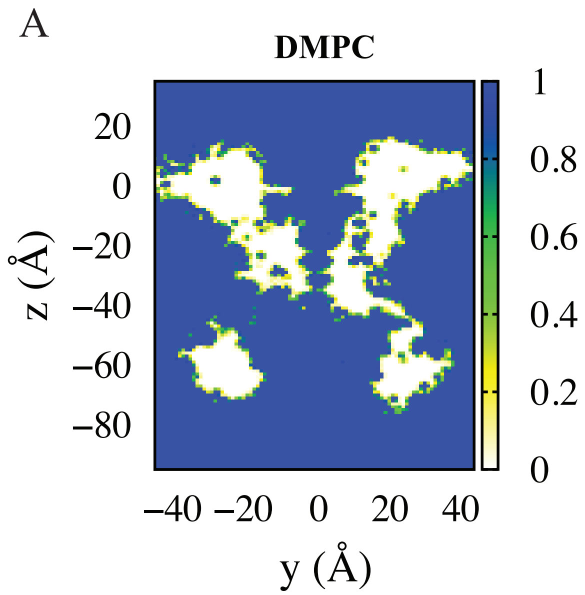


**Figure S5**: **Water density**, illustrated based on the maximum values of water occupancy, is depicted in and around the transporter, as observed within a 1 thick cross-section of the simulation box during the last 5 $\mu$s of both DMPC (A) and POPE (B) simulations.

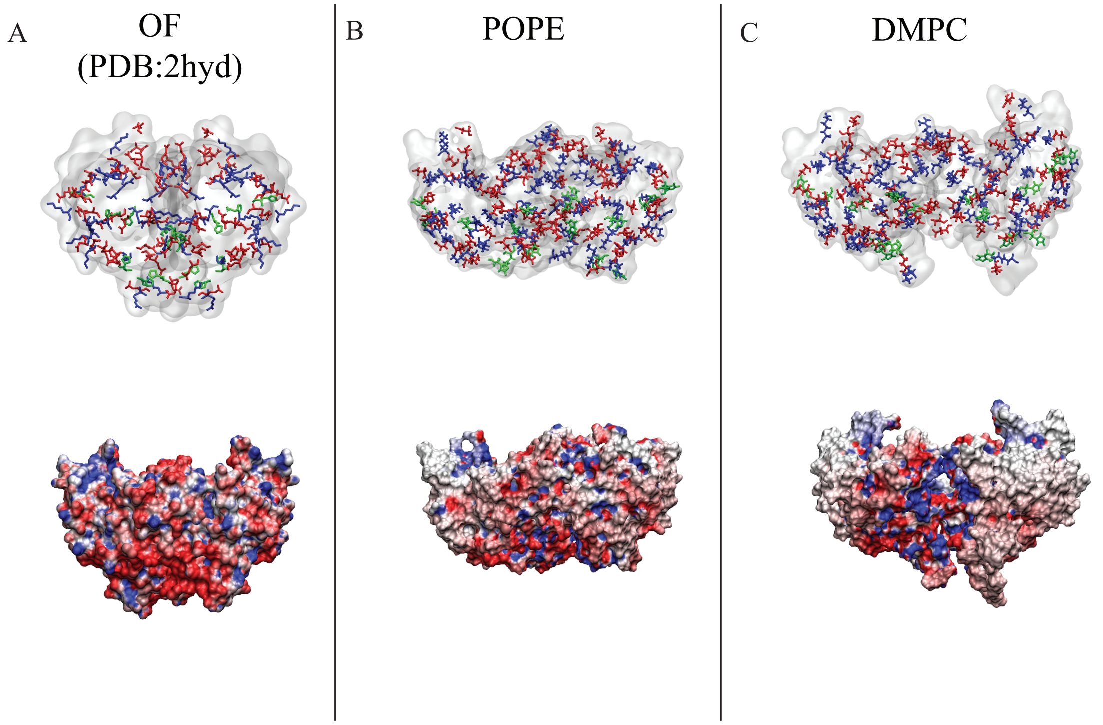


**Figure S6**: **NBD-NBD Interface.** (A, B, C) The top arrow illustrates the charged distribution of positively charged (blue), negatively charged (red), and histidine residues (green) on the dimeric interface. Similarly, the bottom arrow in panels A, B, and C indicates the surface charge distribution for the NBD interface with blue and red denoting positive and negative charges, respectively. In panel A, the snapshot is based on the initial OF conformation (PDB ID: 2HYD). In panels B and C, the surface charge distribution is illustrated for the final conformation of NBDs (10 $\mu$s) in the presence of POPE and DMPC lipid compositions, respectively.

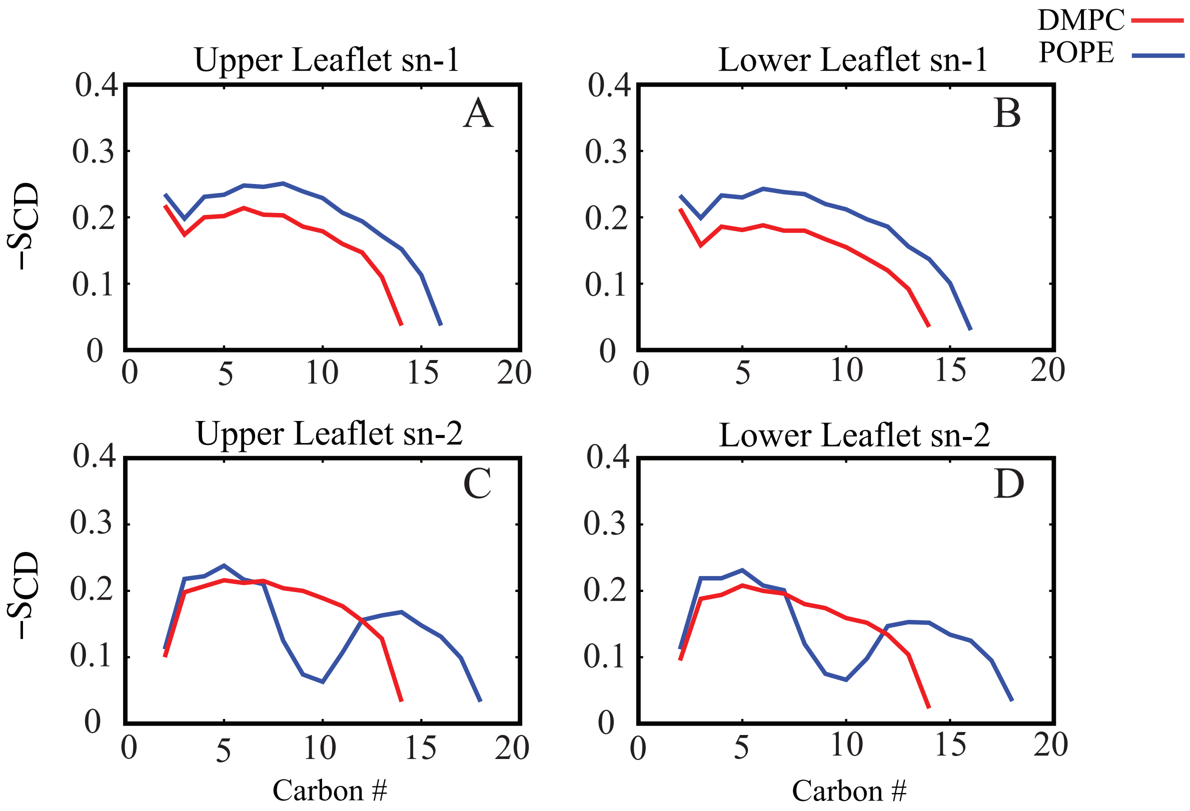


**Figure S7**: **Deuterium Order parameter (SCD) analysis in apo Sav1866 simulations.** Deuterium order parameter (SCD) estimates provide insights into the lipid dynamics within the apo Sav1866 simulation trajectories. The left and right panels illustrate the lipid order parameters associated with the upper (A and C) and lower (B and D) leaflets, respectively. In the color scheme, red represents the DMPC system, while blue represents the POPE system. In the upper and lower leaflets (A and B), the SCDs for chains without double bonds in POPE lipids are notably higher than those in DMPC lipids at 310 K. This suggests a higher degree of lipid ordering in the POPE system, particularly in saturated chains. This enhanced ordering is indicative of increased lipid rigidity and packing in the absence of double bonds. However, the story shifts when considering chains containing a double bond in POPE. In this scenario, the SCD values are lower, especially in carbon numbers around the double bond. This lower SCD indicates increased lipid fluidity and flexibility in the vicinity of the double bond. Therefore, the SCD analysis collectively reveals that while both DMPC and POPE exhibit ordered lipid structures, the introduction of double bonds in POPE disrupts the uniform ordering observed in DMPC, leading to localized areas of enhanced fluidity around the double bond. This distinction in lipid dynamics highlights the sensitivity of Sav1866 to the specific lipid environment and emphasizes the importance of lipid composition in modulating membrane-protein interactions.

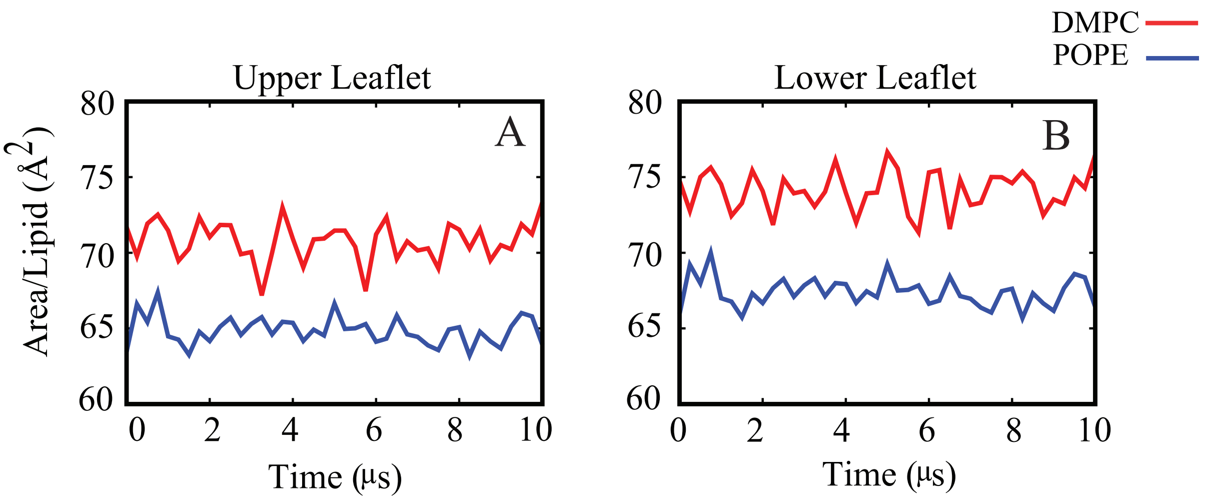


**Figure S8: Analysis of Area per Lipid (APL) in upper and lower leaflets of POPE and DMPC lipids.** Area per lipid (APL) is a fundamental property crucial for understanding membrane characteristics. In the presented analysis, the APL for upper and lower leaflets is shown for POPE and DMPC lipid types in the left and right panels, respectively. DMPC consistently exhibits a greater APL in both leaflets. Specifically, the APL for DMPC initially decreases and stabilizes for the remainder of the simulation, indicating equilibration step. In contrast, for POPE, APL increases initially and stabilizes afterward. This divergence in behavior between DMPC and POPE suggests distinct lipid dynamics and packing arrangements.

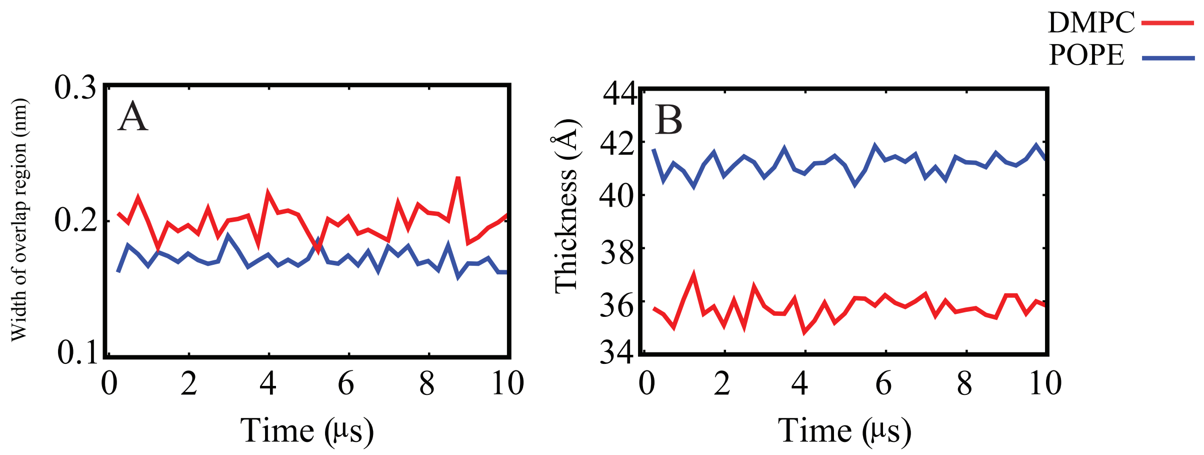


**Figure S9: Analysis of lipid interdigitation and membrane thickness.** The width of the overlap region, as illustrated in Panel (A), serves as an indicator of interdigitation, a phenomenon where the acyl tails of both leaflets overlap at the membrane interface. The comparison between POPE and DMPC lipids reveals a slight difference in the width of the overlap region. Interestingly, DMPC exhibits a higher value for the width of the overlap region compared to POPE, suggesting a potentially greater degree of interdigitation in DMPC. Panel (B) presents the average membrane thickness for DMPC and POPE lipid compositions. Membrane thickness is a complex parameter influenced by factors such as acyl chain length, degree of unsaturation, tilt angle, and lipid composition. Notably, membrane thickness varies between PE and PC lipids. PE lipids consistently exhibit greater thickness compared to PC lipids, reflecting the impact of different headgroup compositions on membrane structure. The greater thickness observed in PE lipids of POPE suggests a more extended and potentially looser packing arrangement compared to the more compact structure of PC lipids in DMPC.

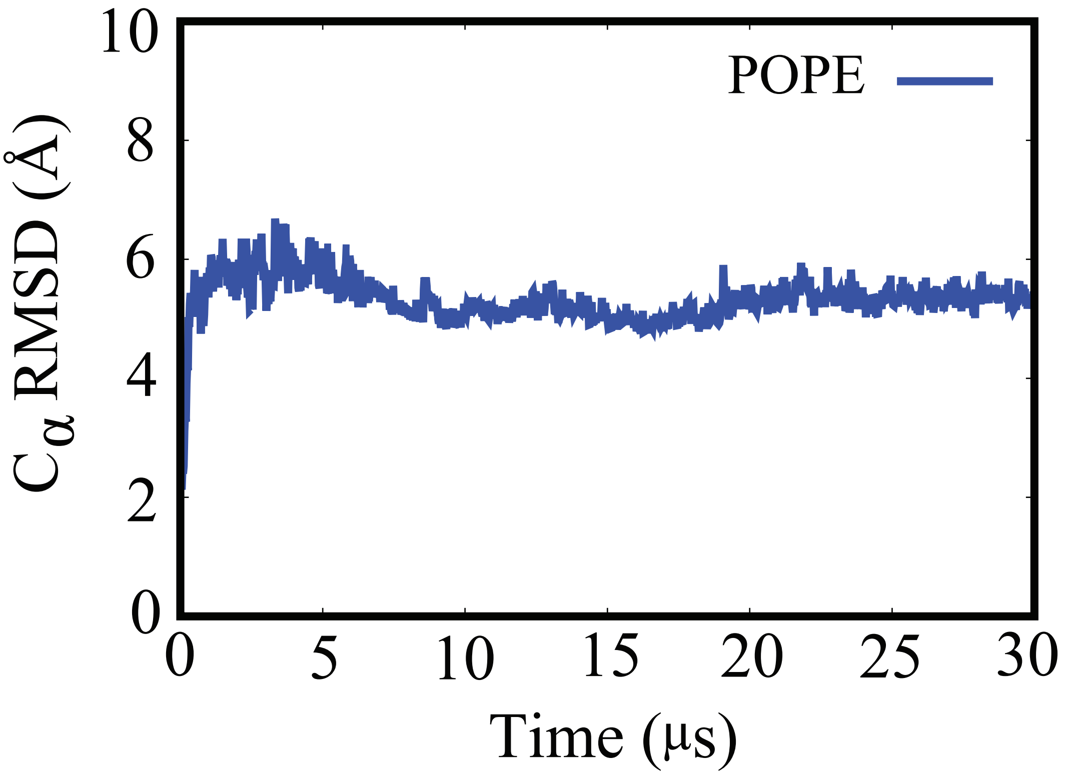


**Figure S10**: **Cα RMSD for 30 $\mu$s simulation.** Time series of root mean square deviation (RMSD) of Cα atoms of the protein associated with apo protein in POPE with respect to the initial model for 30 $\mu$s.

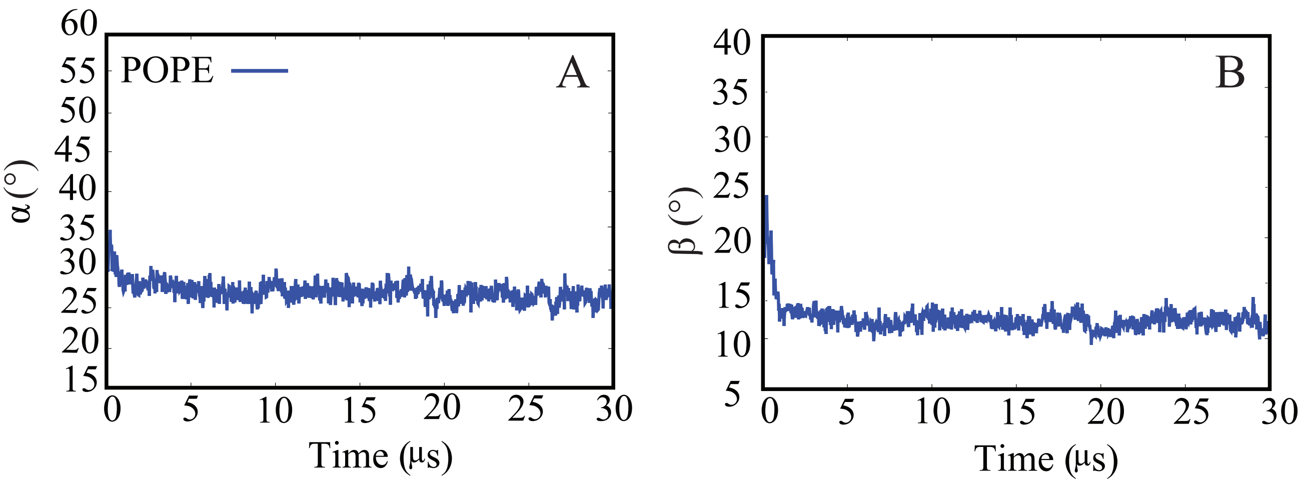


**Figure S11**: $\alpha$ **and** $\beta$ **angle for 30** $\mu$**s simulation.** (A) Time series of the $\alpha$ (cytoplasmic gate) and (B) $\beta$ angle (periplasmic gate) of protein in POPE for 30 $\mu$s.
